# Productive mRNA Chromatin Escape is Promoted by PRMT5 Methylation of SNRPB

**DOI:** 10.1101/2024.08.09.607355

**Authors:** Joseph D. DeAngelo, Maxim I. Maron, Jacob S. Roth, Aliza M. Silverstein, Varun Gupta, Stephanie Stransky, Joel Basken, Joey Azofeifa, Simone Sidoli, Matthew J. Gamble, David Shechter

## Abstract

Protein Arginine Methyltransferase 5 (PRMT5) regulates RNA splicing and transcription by symmetric dimethylation of arginine residues (Rme2s/SDMA) in many RNA binding proteins. However, the mechanism by which PRMT5 couples splicing to transcriptional output is unknown. Here, we demonstrate that a major function of PRMT5 activity is to promote chromatin escape of a novel, large class of mRNAs that we term Genomically Retained Incompletely Processed Polyadenylated Transcripts (GRIPPs). Using nascent and total transcriptomics, spike-in controlled fractionated cell transcriptomics, and total and fractionated cell proteomics, we show that PRMT5 inhibition and knockdown of the PRMT5 SNRP (Sm protein) adapter protein pICln (CLNS1A) —but not type I PRMT inhibition—leads to gross detention of mRNA, SNRPB, and SNRPD3 proteins on chromatin. Compared to most transcripts, these chromatin-trapped polyadenylated RNA transcripts have more introns, are spliced slower, and are enriched in detained introns. Using a combination of PRMT5 inhibition and inducible isogenic wildtype and arginine-mutant SNRPB, we show that arginine methylation of these snRNPs is critical for mediating their homeostatic chromatin and RNA interactions. Overall, we conclude that a major role for PRMT5 is in controlling transcript processing and splicing completion to promote chromatin escape and subsequent nuclear export.

## Introduction

Protein Arginine Methyltransferases (PRMTs) catalyze the methylation of thousands of arginines in hundreds of proteins.^1^ This post-translational modification (PTM) modulates various cellular processes, including RNA processing, transcription, and signal transduction. Based on the nature of the arginine methylation they catalyze, PRMTs are classified into three types: Type I PRMTs (PRMT1, 2, 3, 4, 6, and 8) catalyze asymmetric dimethylation (Rme2a); Type II PRMTs (PRMT5 and 9) catalyze symmetric dimethylation (Rme2s); and a single Type III PRMT (PRMT7) solely catalyzes monomethylation (Rme1).^2,3^ PRMT5 in complex with its obligate cofactor MEP50 is the predominant enzyme responsible for symmetric arginine di-methylation; it has been implicated in various biological processes and diseases, particularly in cancer.

Many PRMT5 studies focused on how PRMT5-mediated arginine methylation affects the assembly and function of the spliceosome via its methylation of three Smith antigen (Sm) proteins: SNRPB (SmB/SmB’), SNRPD3 (SmD3), and SNRPD1 (SmD1).^4-6^ These proteins assemble with four non-methylated Sm proteins into a heptameric ring onto the U1, U2, U4, and U5 splicesomal snRNAs; this assembly protects and organizes the catalytic snRNAs into snRNPs (small nuclear ribonuclear proteins) that are critical for spliceosome function.^6-8^ While inhibition or knockdown of PRMT5 and its cofactors disrupts some RNA splicing, most splicing occurs normally^.9,10^ As measured by total mRNA-seq, both Type I and Type II PRMT inhibition significantly alters gene expression.^1,8^,11 However, the molecular mechanisms by which these global changes occur remain elusive.

Uniquely among PRMTs, PRMT5 has substrate adaptor proteins that recruit specific substrates to enhance their methylation efficiency. pICln contains an Sm protein fold and specifically interacts with the spliceosome proteins SNRPB, SNRPD1 and SNRPD3 to recruit them to PRMT5 and thereby enhance their methylation^.4,10^,12 CoPR5 interacts with PRMT5 and histones H3 and H4.^13^ RIOK1 enhances methylation of nucleolin and the ribosomal protein RPS10.^14,15^

RNA-chromatin interactions are increasingly recognized as vital regulatory mechanisms in the control of gene expression.^16-18^ Poly-adenylated mRNAs, long non-coding RNAs (lncRNAs), and small nuclear RNAs (snRNAs) all can associate with chromatin, in part to influence transcriptional outcomes. For instance, lncRNAs act as scaffolds to recruit chromatin-modifying complexes to specific genomic loci, thereby altering the local chromatin state, higher order chromatin structures, and gene expression.^18^ Similarly, the association of mRNA with chromatin is essential for its proper processing and export from the nucleus^.17,19^ One example of aberrant chromatin-RNA interactions is R-loop (DNA-RNA hybrid) formation, which can induce DNA damage and genomic instability, impacting transcription and higher-order chromatin structures.^20^ We and others previously showed that PRMT5 regulates specific intron detention on chromatin.^8,21^

Despite these advances, the precise molecular mechanisms of gene expression and RNA splicing regulation by PRMT5 remain enigmatic. Here, we investigate mechanisms by which PRMT5 and its adaptor proteins regulate mRNA processing and escape from chromatin. Using small molecule inhibitors of both Type I PRMTs and PRMT5, CRISPR interference (CRISPRi) of PRMT5 and its adaptors, as well as spike-in normalized mRNA transcriptomics, nascent transcriptomics, and proteomics, we reveal surprising roles for arginine methylation in maintaining RNA homeostasis and chromatin dynamics. The loss of arginine methylation on Sm proteins, particularly SNRPB, leads to snRNP and polyadenylated mRNA on chromatin. By fractionating cells and using spike-in normalization approaches, we show that the many studies with total cell mRNA are likely misrepresentative of actual productive changes. Indeed, we observe that most PRMT5 inhibition and PRMT5 and cofactor genetic knockdown mRNA and splicing changes are due to transcripts that remain trapped on chromatin as the result of loss of methylation of spliceosomal snRNP components like SNRPB. Analysis of cytoplasmic transcriptome and proteome consequences reveal that modest productive consequences occur in response to PRMT5 activity loss. Our results demonstrate that PRMT5 plays a critical role in ensuring that transcripts are correctly processed and released from chromatin, thereby maintaining RNA homeostasis and likely preventing genomic instability. Overall, our study highlights a critical role for PRMT5 in post-transcriptional processing of mRNA and productive escape from chromatin and the nucleus.

## Results

### PRMT inhibition results in temporal transcriptome changes

PRMTs catalyze the post-transcriptional methylation of the terminal peptidyl guanidino nitrogens of arginine (**Figure 1a**).^22^ To gain further understanding of the transcriptional consequences of arginine methylation inhibition, we performed poly(A)-RNA sequencing on A549 lung adenocarcinoma cells treated with a time course of DMSO, 1 μM GSK591 (highly selective PRMT5 inhibitor), 1 μM MS023 (Type I PRMT specific inhibitor), or a combination of 1 μM GSK591 and 1 μM MS023 at 2, 4, and 7 days (**Supplemental Table S1**).^8,23,24^ Excessive cell toxicity precluded analysis of 7 day cotreatment data (*data not shown*).

**Figure 1.**
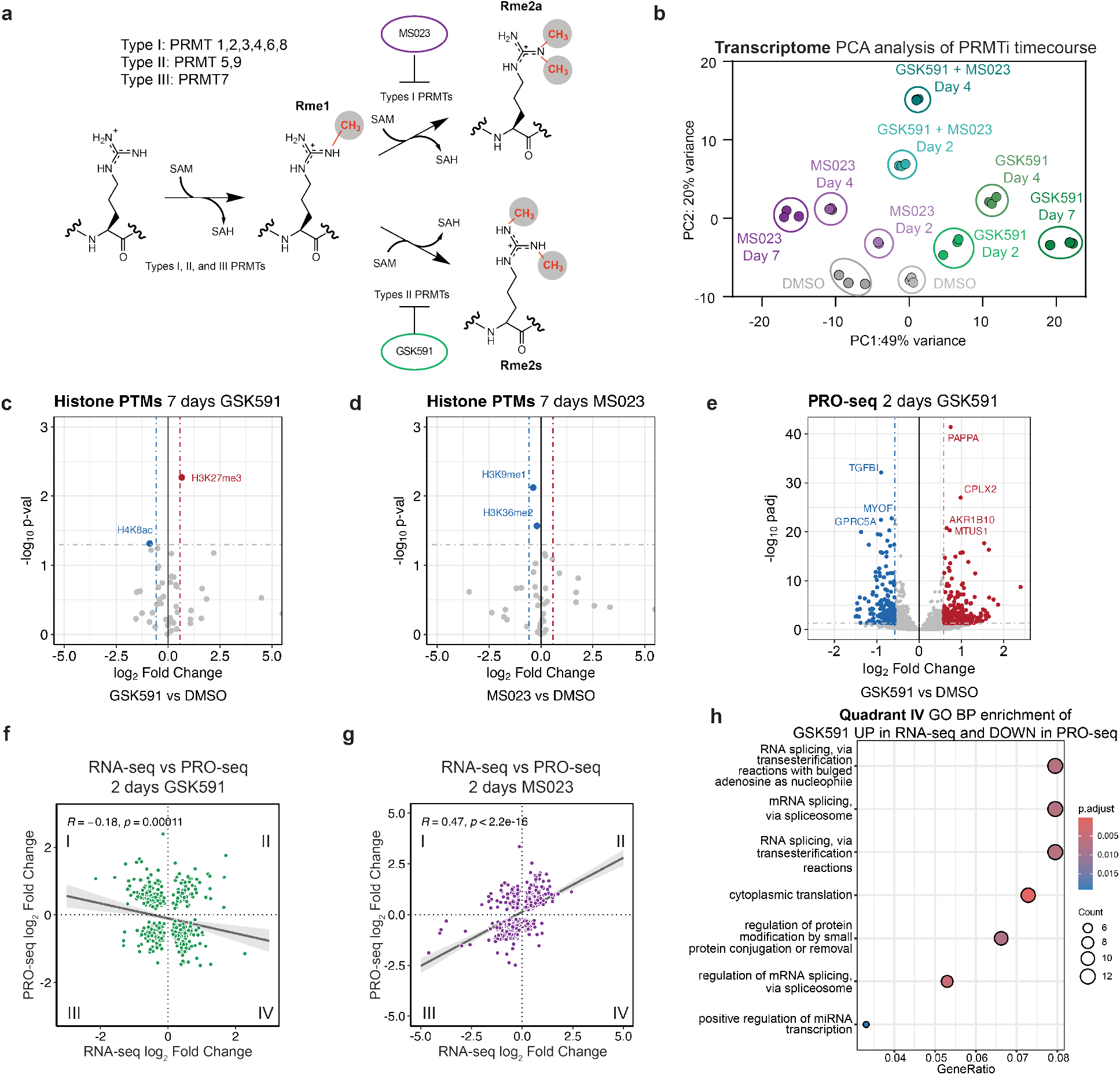
PRMT5 inhibition results in gross transcriptome rearrangements. **a)** Overview of arginine methyltransferases and their methylation reactions. **b)** PCA clustering analysis for an RNA-seq time course of Type I (MS023) and Type II (GSK591) inhibition. **c-d)** Volcano plots of histone PTM proteomics following 7 days of either GSK591 or MS023 treatment. Horizontal lines correspond to a 0.05 p_adj_ cutoff and vertical lines to a 0.58 log_2_ Fold Change. **e)** Volcano plot of nascent Pro-seq after two days of GSK591 treatment, relative to DMSO control. Horizontal lines correspond to a 0.05 p_adj_ cutoff and vertical lines to a 0.58 log_2_ Fold Change. **f-g)** Linear correlations of transcripts significantly called (p_adj_<0.05) in both RNA-seq and PRO-seq experiments for either two days of GSK591 or MS023 treatment. **h)** Dot plot of biological function gene ontology for transcripts upregulated two days GSK591 RNA-seq and downregulated in Pro-seq (Quadrant IV of Figure 1h). Size of the dots corresponds to the number of genes in each category and color is representative of the group’s p-value.

A Principal Component Analysis (PCA) was performed to visualize the transcriptome variance (**Figure 1b**). Principal Component 1 (PC1) showed a temporally dependent variance for both the inhibition of Type I and of Type II PRMTs. This opposite variance on PC1 likely represents the main consequences of Type I and Type II PRMT inhibition. Distance along PC1 increased with time of PRMT inhibition, while PC2 was almost entirely driven by cotreatment with both PRMT inhibitors. Of note, despite transcriptome changes at least as early as two days post-PRMT5 inhibition, growth rate and phenotypic changes associated with PRMT5 inhibition do not occur until later treatment times (**Supplemental Figure S1a**).^1^

To understand the molecular mechanisms driving this transcriptional regulation, we tested how PRMT inhibition influences global histone modifications. We performed histone PTM mass spectrometry analysis on cells treated with either GSK591 or MS023 (**Figure 1c-d**). Surprisingly, despite the large transcriptome changes we observed after 7 days of treatment, there were very few significant changes in histone PTMs with either inhibitor; we also did not observe any histone arginine methylation. The only robust change observed after GSK591 treatment was a modest but significant increase in H3K27me3 abundance, consistent with a previous report linking PRMT5 inhibition with transcriptional repression.^25^ However, these minor changes are unlikely to explain the gross transcriptional changes seen upon PRMT inhibition.

To gain more insight into PRMT transcriptional mechanisms, we further probed the transcriptomic consequences of PRMT5 inhibition. Focusing on the two-day GSK591 time point mRNA-seq, we observed over 1500 significantly altered transcripts, notably prior to any loss of proliferation (**Supplementary Figure S1a-b**). To test if nascent transcription was influenced by PRMT inhibition, we used Precision Nuclear Run-on sequencing (PRO-seq) through a matched time course of enzyme inhibition (**Figure 1e and Supplemental Table S2**, GSK591 2 days; *not shown*: 15 min, 90 min, 3 hours, 2 days, 4 days, and 7 days of both GSK591 and MS023 treatment).^26^ PRO-seq revealed 898 and 509 genes with significantly altered expression following Type I or II PRMT inhibition, respectively (p_adj_<0.05). To understand the interplay of the nascent and mature transcriptome, we intersected significantly altered transcripts between PRO-seq and RNA-seq. Remarkably, there was no correlation between nascent transcription and bulk RNA after two days of Type II PRMT inhibition (**Figure 1f**). This contrasts with a similar comparison of Type I PRMT inhibition with MS023, which resulted in a significant positive correlation (R=0.47, p<2.2×10^−16^) (**Figure 1g**). To determine which gene categories are differentially found in in the nascent and mRNA-seq, we performed gene ontology (GO) analysis on genes upregulated in RNA-seq but downregulated in PRO-seq (lower right quadrant—Quadrant IV—of Figure 1h). Interestingly, this revealed an enrichment in genes involved in RNA homeostasis, including ncRNA processing, RNA splicing, and rRNA regulation (**Figure 1h**). This contrasts with the other three quadrants in which there was no significant enrichment of RNA processing terms (**Supplementary Figure S1c-f**). We conclude that, while Type I PRMTs predominantly regulate nascent transcription, PRMT5 regulates both nascent transcription and post-transcriptional processing.

### PRMT5 adaptor knockdown reveals a PRMT5-pICln-SNRP axis for splicing regulation

To further test the changes in chromatin upon PRMT5 inhibition, we performed immunoblots of acid extracted chromatin from a time course of treated cells (**Figure 2a**).^27^ The major Rme2s-containing protein was observed at approximately 25 kDa, inconsistent with that of histones, but likely corresponding to SNRPB (SmB/SmB’). Furthermore, we observed that both Sm proteins SNRPB and SNRPD3 accumulated on chromatin over GSK591-treatment time, corresponding to their loss of symmetric dimethyl arginine methylation. Consistent with a histone-PTM independent mechanism of transcriptional regulation, only a modest increase in H3K27me3 was observed after 7 days of PRMT5 inhibition; H3K27me3 was not increased after 2 days of treatment (**Figure 2a**). As Sm protein presence in chromatin acid extractions was previously undescribed, we further tested their enrichment by reversed-phase HPLC chromatography (**Supplementary Figure S2a**). Upon subsequent immunoblot analysis, we observed SNRPB in the pooled H2A/H2B/H4 containing fractions (**Supplementary Figure S2b**).

**Figure 2.**
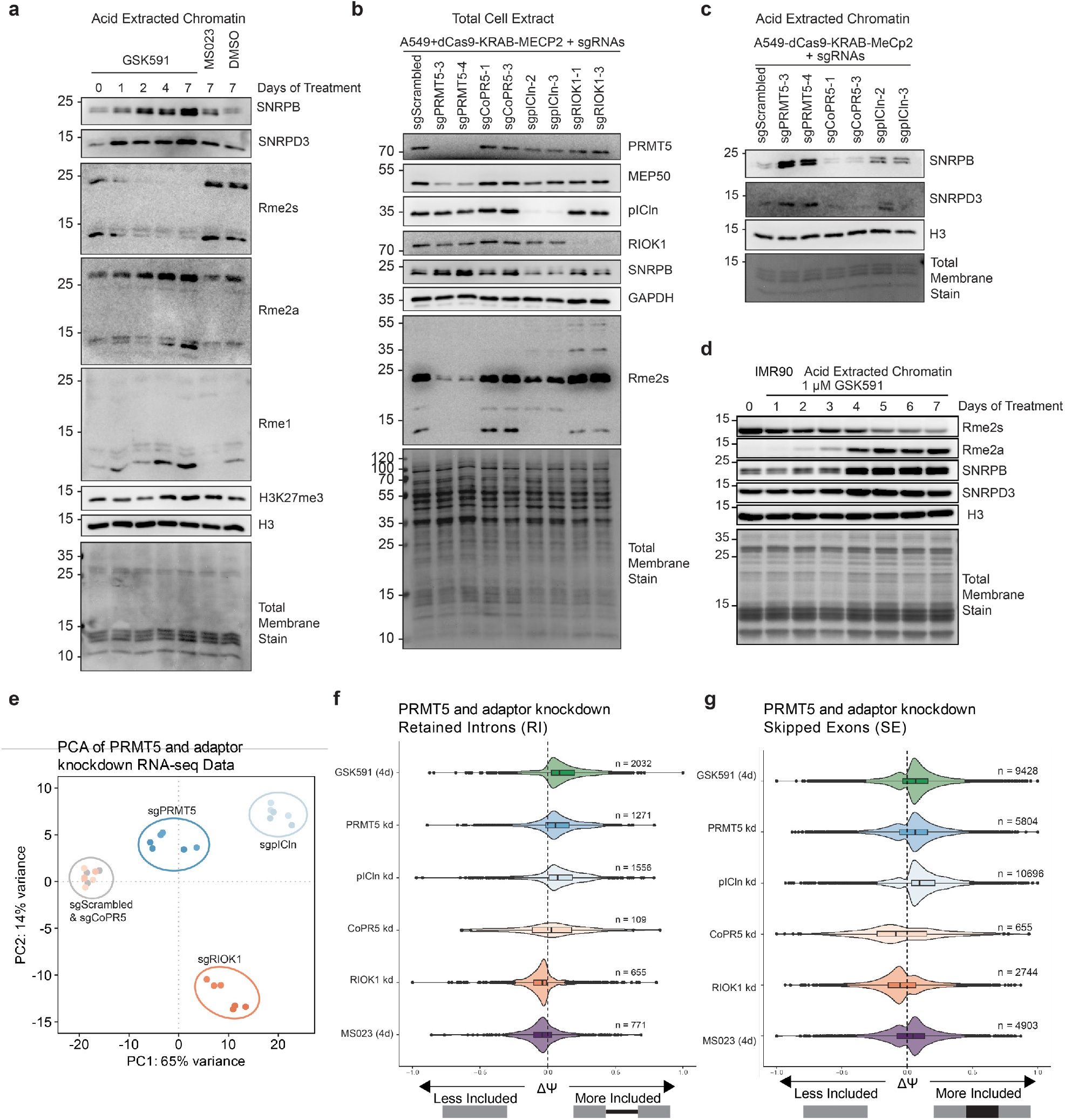
PRMT5-pICln dependent methylation of SNRPB is required for chromatin accumulation. **a)** Immunoblot of acid extracted chromatin over a time course of PRMT5 inhibition by GSK591. DMSO and MS023 controls are also present. **b)** Immunoblot of total cell extract of A549_dCas9-KRAB-MECP2 cells transduced with various sgRNAs. **c)** Immunoblot analysis of acid extracted chromatin of A549_dCas9-KRAB-MECP2 cells transduced with various sgRNAs. **d)** Immunoblot of IMR90-hTert cells over a time course of PRMT5 inhibition. **e)** PCA analysis of RNA-sequencing of PRMT5 and adaptor protein knockdowns. **f)** Comparison of ΔΨ (Delta PSI / Percent Spliced In) of retained introns (RI) following PRMT5 and adaptor protein knockdowns. **g)** Comparison of ΔΨ (Delta PSI) of skipped exons (SE) following PRMT5 and adaptor protein knockdowns.

This accumulation of non-methylated Sm proteins prompted us to consider whether Sm protein methylation, rather than histone arginine methylation, could be a key mechanism of PRMT5 transcriptional regulation. Therefore, we sought to clarify which PRMT5 substrates were responsible for the observed transcriptional changes and Sm chromatin accumulation. To accomplish this, we used CRISPRi to knockdown PRMT5 and three of its substrate adaptors: pICln, CoPR5, and RIOK1.^10,13,14,28-32^ We observed substantial knockdown of each target protein via immunoblots and RT-qPCR (**Figure 2b** and *not shown*). Consistent with our previous work, loss of PRMT5 resulted in a loss of its obligate complex member MEP50; however, knockdown of the other PRMT5 adaptors did not have a major effect on either PRMT5 or MEP50 protein levels (**Figure 2b)**.^33,34^

To determine cellular phenotypic consequences of the adaptor knockdowns, we performed growth assays and cell cycle analyses (**Supplemental Figure S2d-g**). Compared to GSK591-treatment, the PRMT5 knockdowns exhibited similar growth inhibition as well a G1 cell cycle arrest (**Supplemental Figure S2h**); this is consistent with observations in previous studies.^35,36^ We observed similar growth inhibitions in both pICln and RIOK1 knockdowns but not in the CoPR5 knockdowns. While pICln and RIOK1 knockdowns had similar G1 arrest to the PRMT5 knockdown, they also exhibited an increase in G2/M phase and a decrease in S phase, suggesting both PRMT5-mediated and independent phenotypes.

In total lysate immunoblots, we observed a modest decrease in SNRPB protein in PRMT5 knockdown and a more pronounced loss in the pICln knockdown (**Figure 2b**). This is consistent with a report of Sm lysosomal degradation upon pICln loss.^37^ We also observe a significant decrease in methylated SNRPB and SNRPD3 in both the PRMT5 and pICln knockdowns.

To understand the mechanisms mediating Sm-chromatin accumulation, we performed immunoblot analysis of acid extracted chromatin from the PRMT5 and adapter knockdowns (**Figure 2c**). We observed SNRPB and SNRPD3 accumulation on chromatin only upon pICln and PRMT5 knockdown. Overall, we concluded that SNRP arginine methylation via a PRMT5-pICln axis drives SNRP accumulation on chromatin.

We further tested if SNRPB and SNRPD3 chromatin accumulation upon PRMT5 inhibition is a general phenomenon in mammalian cells. We treated both IMR90-hTERT cells, immortalized fibroblasts from normal human lung tissue, and NSC-34 cells, a mouse motor-neuron like cell line, with either Type I or Type II PRMT inhibition **(Figure 2d** and **Supplemental Figure S2c)**. In both cell lines, we observed SNRPB and SNRPD3 accumulation with concomitant loss of Rme2s, demonstrating that SNRP-chromatin retention is a widespread consequence of PRMT5 inhibition.

Next, we studied the effects of adaptor knockdowns on the transcriptome. We performed bulk poly(A) RNA-sequencing on two individual knockdowns for PRMT5 and each of the three substrate adaptors, compared to a non-targeting scrambled sgRNA control (**Supplemental Table S1**). Principal component analysis and a z-score heatmap demonstrated clear distinctions between the knockdown of the different adaptor proteins (**Figure 2e** and **Supplemental Figure S2i**). PC1 captured the majority of variance and was separated by PRMT5, pICln and RIOK1 from the scrambled control and CoPR5. As CoPR5 clustered with the scrambled control it had no significant effect on the transcriptome. Variance along PC2 was driven by RIOK1, while PRMT5 and pICln clustered similarly (**Figure 2e**).

As the abundant RNA splicing consequences of PRMT5 inhibition are well established, we examined if any of the adaptor knockdowns would replicate these effects.^8,38^,39 Consistent with its role in enhancing Sm methylation, pICln was the only adaptor to mimic both PRMT5 inhibition and knockdown splicing consequences, such as retained introns (RI) and skipped exons (SE) (**Figure 2f-g**). Notably, the overlap of shared retain introns was significant both by Jaccard Index and Fisher Exact Test (p<0.05), compared to the universe of all expressed introns in A549 cells (**Supplemental Figure S2j-k**). This is consistent with prior work implicating the PRMT5-pICln axis in the regulation of mRNA splicing.^40^

Overall, these studies revealed that knockdown of PRMT5 and its adapters produced unique transcriptomic consequences. Moreover, these studies revealed the role of PRMT5-pICln axis in regulating splicing consequences and concurrent SNRPB and SNRPD3 retention on chromatin.

### PRMT5 regulates SNRPB abundance and subcellular localization

To further understand how PRMT5 regulates transcription, we investigated the proteomic consequence of PRMT5 inhibition. First, we performed total proteomic mass spectrometry on cells treated with GSK591 for two days (**Figure 3a** and **Supplemental Table S3**). While only a small number of proteins were significantly changed in abundance (18 total proteins, p_adj_ < 0.1), we observed that the most upregulated protein was SNRPB; other upregulated proteins included splicing and RNA-processing proteins, including SART1, PRPF4B, and PRPF39. Comparing the small number of proteomic changes with the transcriptome after 2 days of PRMT5 inhibition revealed a moderate positive correlation (R=0.64, p=4.1×10^−9^) (**Figure 3b**).

**Figure 3.**
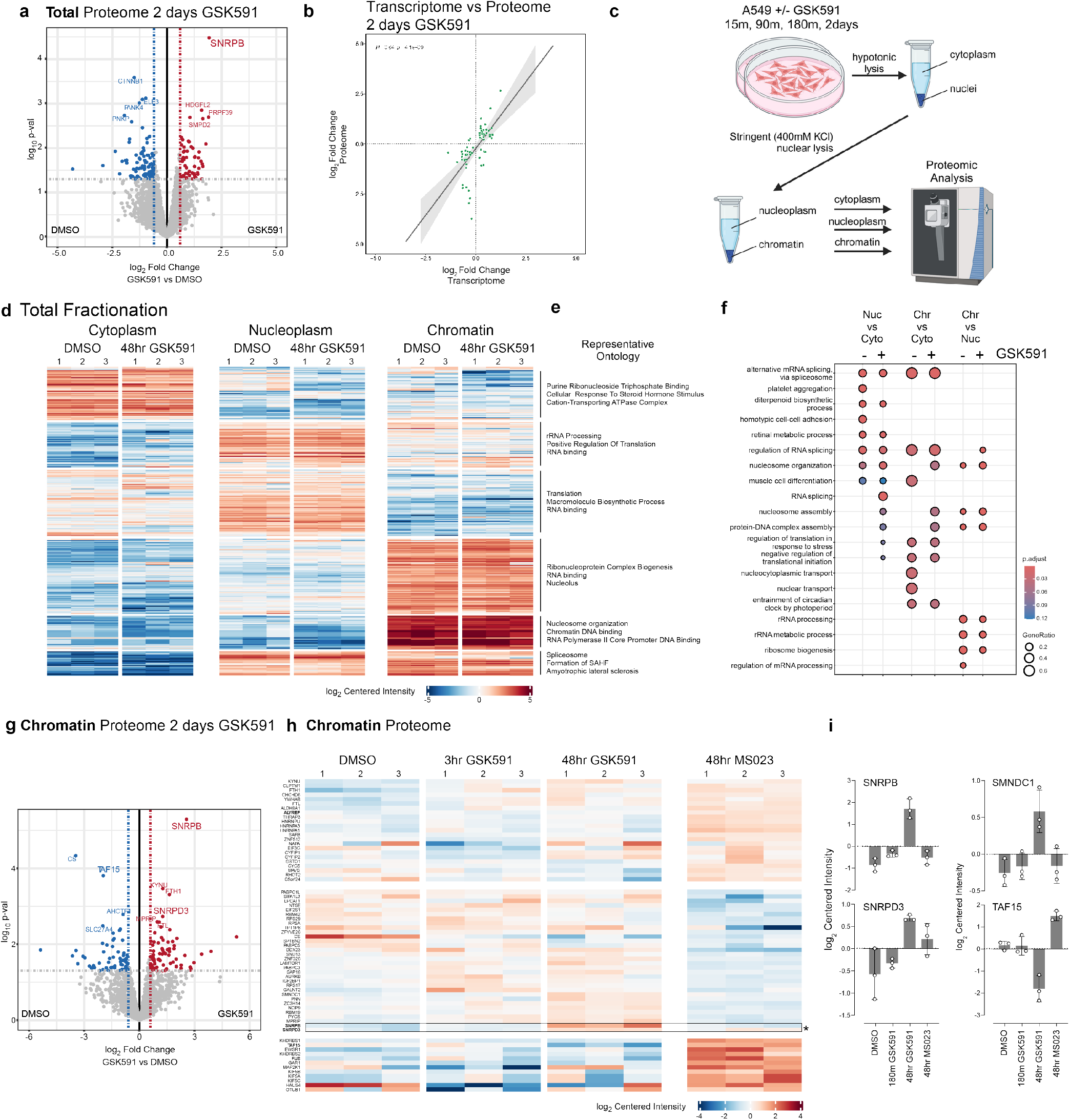
Fractionated proteomics reveal that SNPRB accumulation on chromatin is the major proteomic consequence of PRMT5 inhibition. **a)** Volcano plot of total cell proteome following two days of GSK591 inhibition. Highlighted proteins are those above a p-value>0.05 and an absolute log_2_ fold change >0.58. **b)** Linear correlation of shared genes from total RNA-sequencing and proteomics (with a p_adj_ <0.05 cutoff) following two days of PRMT5 inhibition. **c)** Schematic of cellular fractionation approach. **d-e)** Heatmap and representative ontology for each cellular fraction and treatment. Color corresponds to a log_2_ of centered intensity. **f)** Dot plot highlighting ontology of biological function of proteins in each cellular compartment. Size of dot corresponds to gene ratio and color to the adjusted p-value. **g)** Volcano plot of mass-spec log_2_ Fold change vs log10 p_val_ of the chromatin fraction following GSK591 treatment. **h)** Heatmap comparing changes in the chromatin proteome. Block color is representative of log_2_ centered intensity. **i)** Selected histograms of chromatin associated proteins and their centered intensity across different drug treatments.

As we had initially observed SNRPB and SNRPD3 accumulation on chromatin post-PRMT5 inhibition (**Figure 2a**), we tested changes in the subcellular distribution of the proteome in a Type I PRMT and PRMT5-dependent manner. To perform a fractionated cell proteomic analysis, we developed a stringent cellular fractionation protocol to efficiently separate cytoplasmic, nucleoplasm, and chromatin proteins (**Figure 3c**). We treated cells with PRMT inhibitors GSK591 and MS023 over time (15 min, 90 min, 180 min, and 48 hours, in biological triplicate), isolated protein from the cytoplasmic, nucleoplasm, and chromatin fractions, and subsequently performed mass spectrometry. We then determined the differential abundance of proteins between each fraction and drug treatment condition. To test enrichments and changes between these conditions, we used k-means clustering of the top 340 proteins altered and plotted a heatmap of the centered normalized intensities for the DMSO control and the 48-hour GSK591 treatment (**Figure 3d**; other time points *not shown*). The major clusters clearly separated the cytoplasm, nucleoplasm, and chromatin. Representative GO ontologies for each of the six k-means clusters are indicated; these were consistent with ontologies expected for the subcellular fraction in which they were enriched (**Figure 3e**). For instance, highly enriched proteins on chromatin were represented by nucleosome organization terms. Comparing 48-hour GSK591- and DMSO treated chromatin proteomes revealed gene ontology alteration of proteins related to RNA processing, RNA binding, and splicing (**Figure 3e-f**). This is consistent with the transcriptomic ontology analysis at similar two days of PRMT5 inhibition (**Supplementary Figure 1a**).

Focusing specifically on the chromatin fraction, the most upregulated protein was SNRPB (**Figure 3g-i**); this was consistent with the chromatin immunoblots (**Figure 2a**). SNRPD3, also methylated by PRMT5, is similarly enriched on chromatin. We also observed TAF15 chromatin depletion after PRMT5 inhibition; this was consistent with reports that arginine methylation affects its cellular localization.^41^

### Unmethylated SNRPB chromatin accumulation is transcription-dependent

Having confirmed that SNRPB and SNRPD3 chromatin accumulation is a consequence of PRMT5 inhibition, we aimed to further understand this phenotype. As some prior literature suggested that Sm methylation is involved in UsnRNP assembly via its interaction with SMN^4,5^, we first asked whether methylation of Sm proteins via PRMT is required for their proper assembly into mature snRNPs. To test whether arginine methylation of SNRPB or SNRPD3 affects their assembly into mature snRNPs, we performed a co-immunoprecipitation assay, enriching for the RNA Trimethyl-guanosine (TMG) cap—a marker of mature, fully assembled snRNPs—in either control or PRMT5 inhibited cells, followed by subsequent immunoblotting for Sm proteins (**Figure 4a**).^42,43^ U1-70k and SNRPE were positive controls for the co-immunoprecipitation as they are non-arginine methylated components of mature snRNPs^.44,45^ Strikingly, there was no difference in SNRPB or SNRPD3 levels with PRMT5 inhibition; highlighting that arginine methylation is not necessary for assembly of mature snRNPs. This result is consistent with our and others work that demonstrated Sm proteins lacking arginine in their tails were incorporated into snRNPs and imported into the nucleus^8,46^ and provides further support that PRMT5-mediated methylation of Sm proteins is not required for proper snRNP assembly. We therefore ruled out impaired UsnRNP assembly as the mediator of PRMT5-mediated transcriptional regulation.

**Figure 4.**
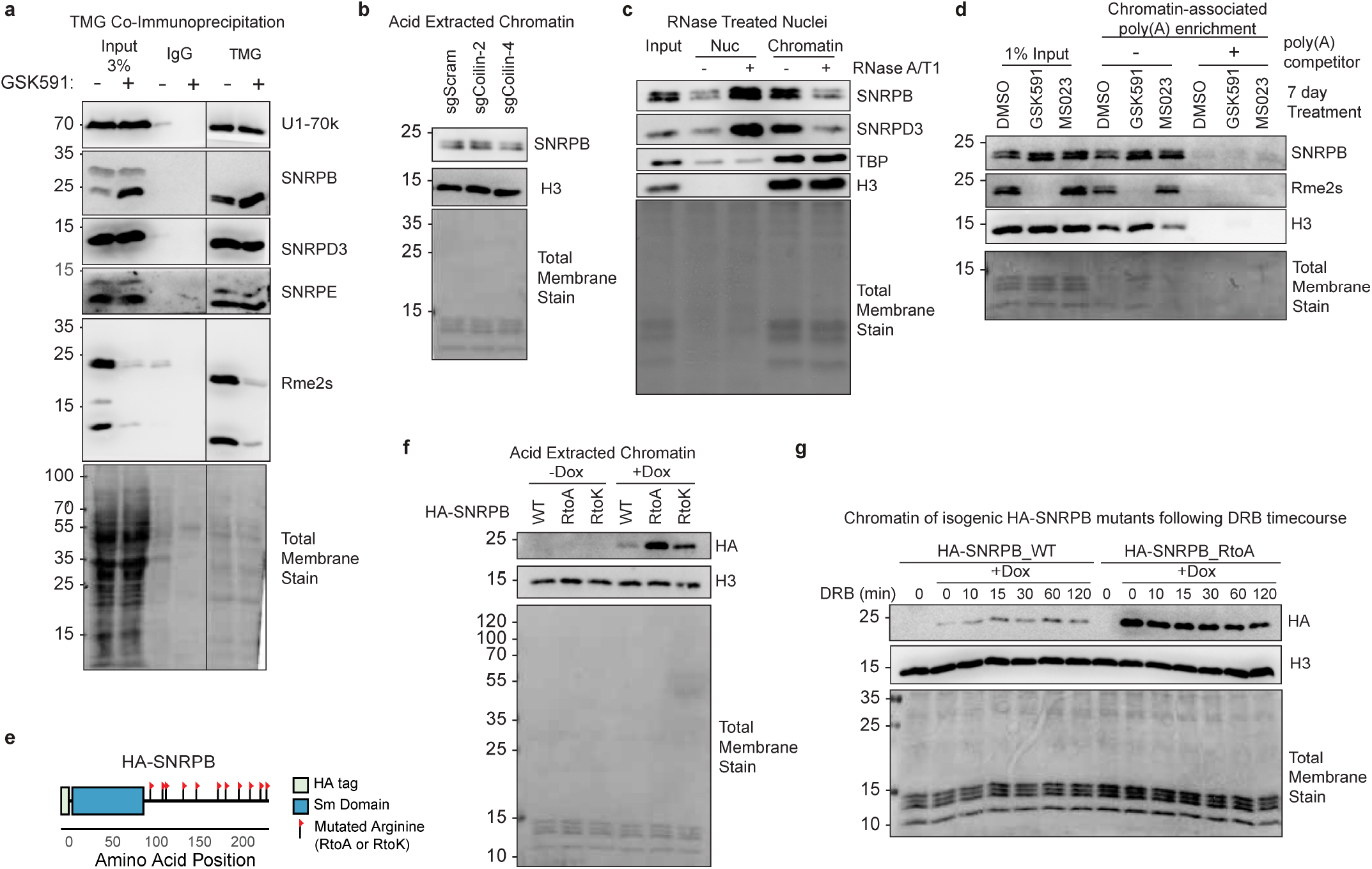
Unmethylated SNRPB chromatin accumulation is dependent on post-transcriptional mRNAs. **a)** Co-immunoprecipitation of TMG followed western blotting for Sm proteins and mature snRNP components. **b)** Immunoblots of acid extracted chromatin following Coilin CRISPRi knockdowns. **c)** Immunoblots of nucleoplasm and chromatin fractions after nuclei treatment with RNase A/T1. **d)** Immunoblot of pulldown with oligodT-linked beads with crosslinked chromatin. A poly(A) competitor was used as a negative control for the pull-down. **e)** Schematic of cloned HA-SNRPB ORF with each flag representing a site of arginine mutation on the C-terminal tail. **f)** Immunoblot of acid extracted chromatin of inducible SNRPB construct over-expression with doxycycline treatment. **g)** Immunoblot of chromatin after inducing SNRPB WT and RtoA expression and a time course of DRB treatment.

Next, we asked if PRMT5-inhibition dependent Sm chromatin accumulation is a consequence of disrupted nuclear Cajal bodies. As Cajal bodies are a site of snRNP assembly and recycling, we asked if loss of the Cajal body organizing protein Coilin—and concomitant impaired recycling of Sm proteins in snRNPs—is responsible for Sm accumulation on chromatin.^47-49^ After CRISPRi knockdown, we observed no significant difference in SNRPB chroma-tin-accumulation compared to a non-targeting control knockdown (**Figure 4b** and **Supplemental Figure S3a**). We therefore concluded that Cajal bodies are not responsible for SNRPB chromatin detention.

To test if intact snRNPs or just Sm proteins were chromatin detained, we performed Northern blotting with snRNA probes. In GSK591-but not MS023-treated cell nucleoplasm and chromatin fractions, these blots revealed an increased amount of U2, U1, U4, U5, and U6 major snRNAs (**Supplemental Figure S3b**). These observations were consistent with intact snRNPs accumulating on chromatin in a PRMT5-inhibited fashion.

As Sm proteins are part of RNA binding complexes, we tested if the increased SNRPB and SNRPD3 chromatin affinity was RNA dependent. We treated cells with GSK591, followed by nuclear extraction, combined RNase A/T1 treatment, and nucleoplasm and chromatin isolation and immunoblotted for Sm proteins; we used the DNA-binding TATA-binding protein (TBP) as a positive control. Upon RNA removal, we observed a clear loss of chromatin associated SNRPB and SNRPD3 as well as a gain of soluble nucleoplasm proteins (**Figure 4c**). As we previously showed that SNRPB was bound to poly(A)-RNA^13^, we next asked whether the chromatin poly(A)-RNA bound Sm proteins with or without Rme2s. Strikingly, after GSK591-treatment and despite the complete absence of Rme2s on SNRPB, its binding to poly(A) transcripts on chromatin was not disrupted (**Figure 4d**).

To test if Sm-chromatin accumulation was directly due to Sm arginine methylation, we created an isogenic system.^8,49^ We engineered a SNRPB overexpression system in which we mutated each arginine in the C-terminal tail to either an alanine (RtoA) or lysine (RtoK) (**Figure 4e**). Each of these constructs consistently expressed at comparable levels following a four-day induction with doxycycline (DOX) (**Supplemental Figure S3c**). To test the propensity of unmethylated SNRPB to accumulate on chromatin, we induced the mutant expression with DOX followed by chromatin acid extractions. We observed an increase in the amount of chromatin-bound RtoA and RtoK compared with wildtype SNPRB (**Figure 4f**). These results are consistent with SNRPB accumulation on chromatin as a direct consequence of its arginine methylation and not a consequence of some other PRMT5-methylated targets.

We next asked if SNRPB chromatin retention was also dependent on RNA Polymerase II (Pol II) function. To test this, we DOX-induced both SNPRB WT and RtoA mutant, followed by a time course of 5’6-dichlorobenzimidazole (DRB), an inhibitor of Pol II pause release.^50^ We observed only a modest decrease in the supra-normal chromatin bound SNRPB RtoA mutants with DRB treatment (**Figure 4g**). Taken together, these data demonstrate that SNRPB chromatin retention is mediated by a loss of arginine methylation on its intrinsically disordered tails and through its interaction with mature, poly(A) RNAs.

### PRMT5 inhibition results in mRNA chromatin detention

As we determined that SNRPB and SNRPD3 accumulated on chromatin in an RNA-dependent manner, we next sought to understand to what extent RNA itself was being detained on chromatin. To accomplish this, we performed a time course of PRMT5 inhibition followed by simultaneous RNA and DNA extractions using TRIzol^51^ and quantified the total amount of RNA relative to DNA. This revealed increasing amounts of total RNA concomitant with duration of PRMT5 inhibition (**Figure 5a**). Of note, the increase in RNA abundance preceded increased cell volume and decreased growth rate that we observe in A549 cells after PRMT5 inhibition (**Supplemental Figure S4a** and **Figure 1c**). To avoid confusing increased RNA with changes in cell size or growth arrest we focused our studies on two days of PRMT5 inhibition.^52^ Next, we determined which cellular compartment was responsible for this RNA increase. To accomplish this, we isolated RNA from cytoplasm, nucleoplasm, and chromatin fractions using an optimized a cellular fractionation protocol (**Figure 5b**).^8,19,53,54^ DNA and protein were simultaneously extracted from the chromatin fraction to both normalize RNA quantities and provide fractionation controls, respectively. Strikingly, this experiment revealed that RNA accumulated specifically within the chromatin fraction. Upon PRMT5 inhibition we found a significant increase in the amount of chromatin-bound RNA (p<0.05) (**Figure 5c-d**). Of note, as there was no notable RNA accumulation on chromatin even after 7 days of Type I inhibition by MS023, this RNA accumulation was specific to PRMT5 inhibition (**Supplemental Figure S4b**).

**Figure 5.**
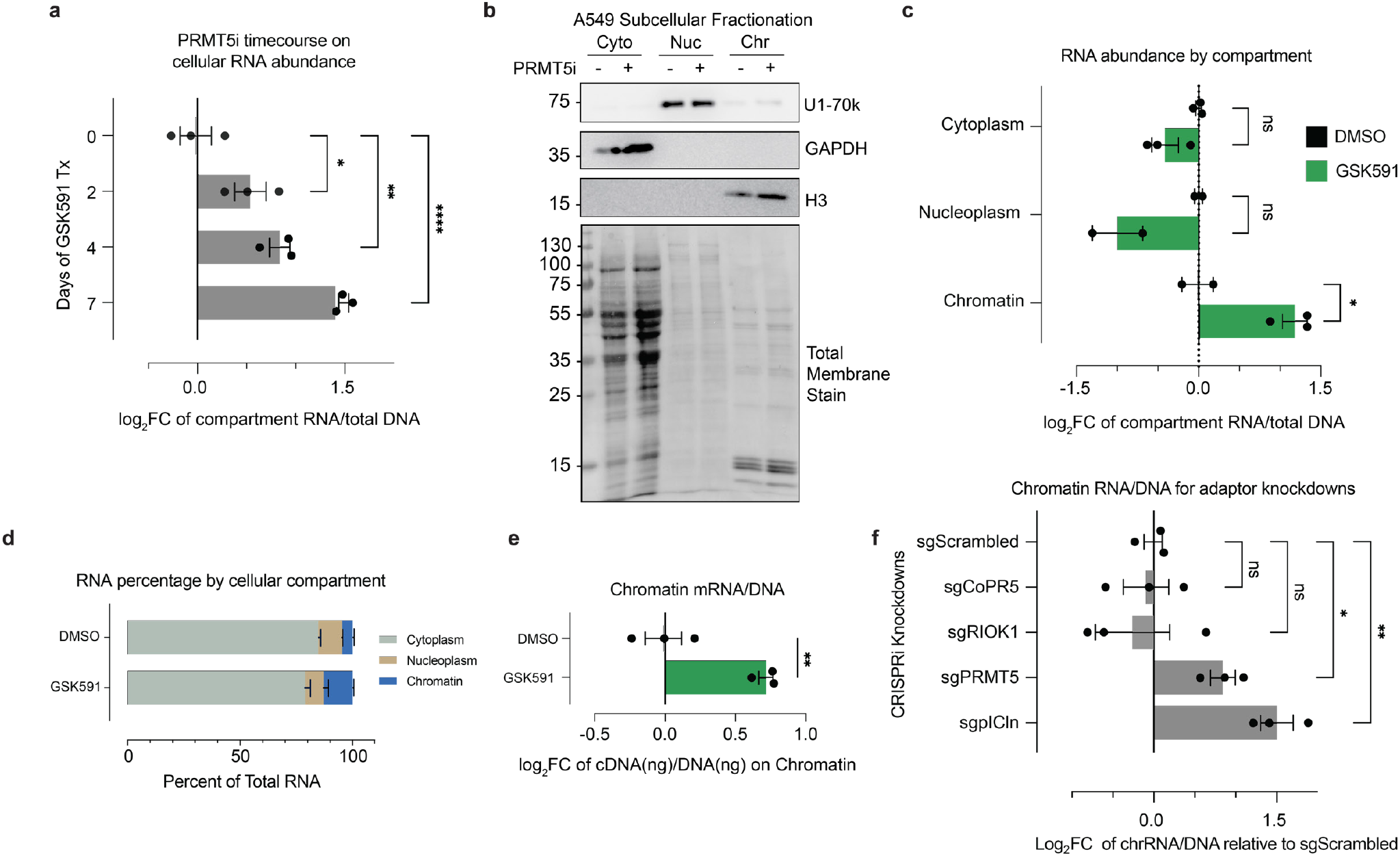
PRMT5 inhibition results in mRNA chromatin detention. **a)** Histograms depicting the amount of total RNA per cell normalized to the amount of DNA. *<0.01, **<0.01, ****<0.0001, n.s. = not significant. **b)** Immunoblots validation of RNA cellular fractionation protocol into cytoplasm, nucleoplasm, and chromatin fractions. **c)** Gross RNA abundance per cellular compartment, normalized to DNA in the chromatin fraction. *<0.01, n.s. = not significant. **d**) Relative abundance of RNA per cellular compartment with GSK591 or DMSO control. **e)** Gross amount of mRNA on chromatin, normalized to DNA. mRNA measured via RNA conversion to cDNA using oligo dT primers. **f)** Relative abundance of RNA on chromatin for PRMT5 and adaptor protein knockdown, normalized to amount of cellular DNA per sample. *<0.01, **<0.01, n.s. = not significant.

We next asked if mature, polyadenylated RNA was trapped on chromatin. Using chromatin-isolated RNA, we performed a reverse transcriptase reaction with oligo-dT priming and generated cDNA. The cDNA from polyadenylated RNAs was then normalized to total DNA per sample (**Figure 5e**). We observed a significant PRMT5-dependent enrichment of mature, polyadenylated RNA transcripts on chromatin.

Finally, to gain insight into which PRMT5 methylation substrates may be responsible for the observed RNA chromatin accumulation, we utilized the PRMT5 adaptor knockdowns. sgRNAs targeting each adaptor protein were transduced into the stable A549 dCas9 cell line and passaged for 7 days post selection. Cells were subsequently fractionated and the chromatin RNA/DNA ratios were measured (**Figure 5f**). The PRMT5 knockdown recapitulated the inhibitor-based observation of increased RNA on chromatin. For the adaptor protein knockdowns, only pICln resulted in increased RNA on chromatin, supporting that this phenotype occurred due to decreased Sm protein arginine methylation.

### PRMT5 is required for polyadenylated mRNA transcript escape from chromatin

Having established that PRMT5-inhibition causes mature, polyadenylated RNAs to accumulate on chromatin, we next sought to identify these transcripts by poly(A) enriched RNA-sequencing. We pretreated cells with two days of PRMT5 inhibition, followed by fractionation into cytoplasm, nucleoplasm, and chromatin (**Figure 6a, Supplemental Figure S5a-b**, and **Supplemental Table S4**). To rigorously detect changes in RNA abundance that may be otherwise missed by standard bioinformatic normalization methods, we used a spike in of *Drosophila melanogaster* S2 cells (5% relative to starting cell number).^55-57^ Importantly, when normalizing the transcriptome to the *Drosophila* RNA, we observe that sequencing of the spiked-in, cross-species RNA was consistent between samples (**Supplemental Figure S5c**).

**Figure 6.**
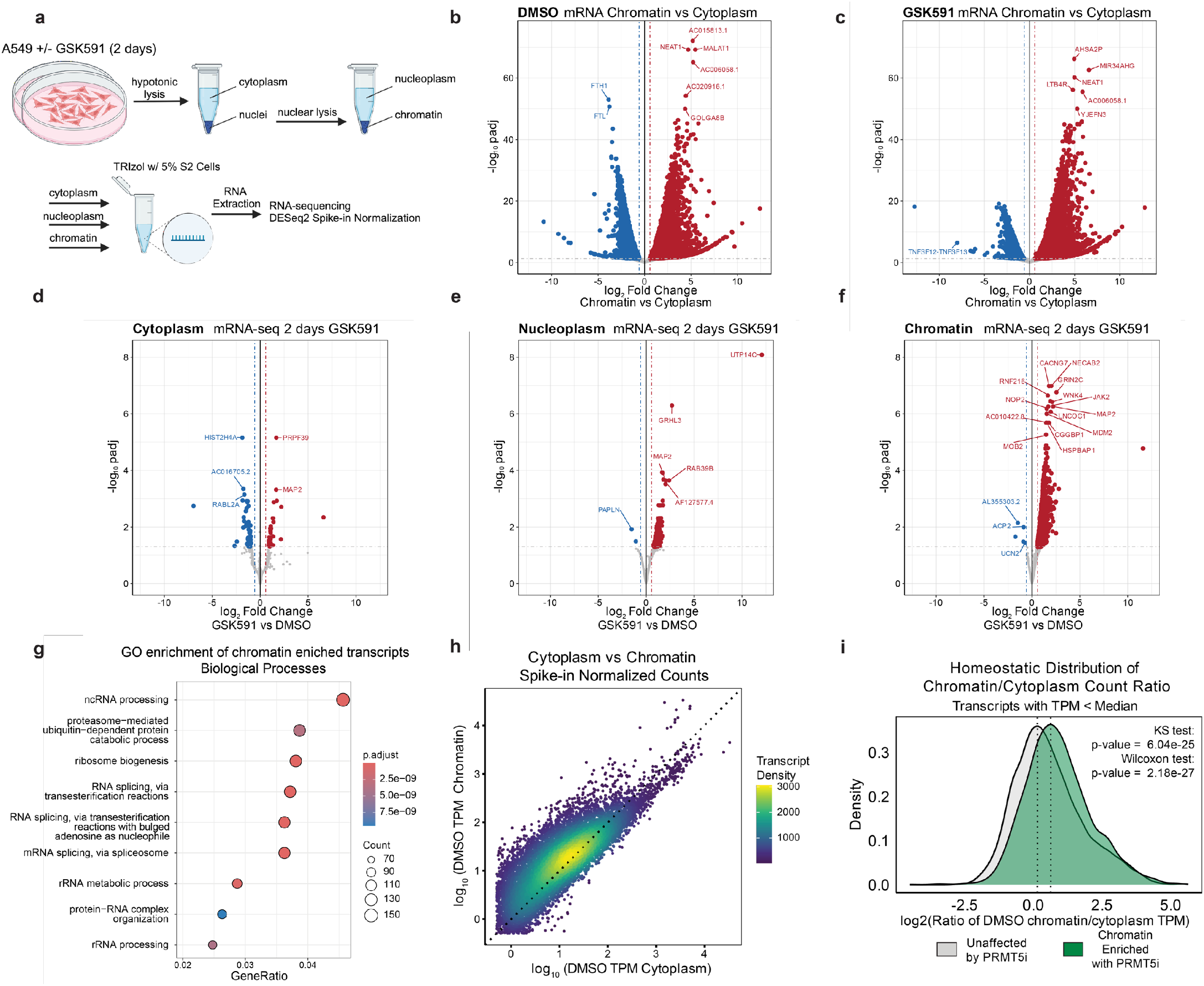
PRMT5 promotes mRNA escape from chromatin. **a)** Schematic of cellular fractionation and RNA-sequencing experiment. **b-c)** Volcano plots of mRNA-sequencing of each treatment comparing cytoplasm to chromatin using S2 spike-in normalization. Red transcripts have a log2FC>0.58 and a p_adj_>0.05. Blue transcripts have a log2FC<-0.58 and a p_adj_>0.05. **d-f)** Volcano plots of mRNA-sequencing of each cellular compartment between conditions using S2 spike-in normalization. Red transcripts have a log2FC>0.58 and a p_adj_>0.05. Blue transcripts have a log2FC<-0.58 and a p_adj_>0.05. **g)** Dot plot of biological function gene ontology for chromatin-enriched transcripts. Dot size is representative to the number of genes per category and color represents the p-adjusted value. **h)** Correlation of the log_10_ average spike-in normalized read counts in TPM for transcripts in the cytoplasm and chromatin compartments for untreated cells (DMSO). Dotted line represents y=x. Transcript color represents the density of points on the plot. **i)** Density plots of the ratio of normalized TPM counts per gene (with TPM < median of all expressed genes) between chromatin and cytoplasm compartments, comparing genes found to be enriched on chromatin upon PRMTi. Kolmogorov-Smirnov (KS) and Wilcoxon ranked sum tests were used to compare distributions.

First, we compared the relative abundance of cytoplasmic to chromatin transcripts with PRMT5 inhibition (**Figure 6b-c**). This revealed a striking enrichment in chromatin-associated RNAs following PRMT5 inhibition. While some transcripts were predominantly cytoplasmic, others were enriched in the chromatin fraction, indicating that they are not being exported into the cytoplasm.

Next, we compared the relative abundance of transcripts between PRMT5-inhibited and control cells. In the cytoplasmic mRNA, we observed few changes, with only 80 significantly altered transcripts (p_adj_ < 0.05) (**Figure 6d**). This is consistent with our findings of minimal changes in the total proteome (**Figure 3a**). In the nucleoplasm, there were 409 significantly altered transcripts, 407 of which were increased in abundance (**Figure 6e**). Remarkably, when examining the chromatin fraction, we observed nearly 4,000 transcripts that were increased in abundance with only a handful being downregulated (**Figure 6f**). Most of the altered polyadenylated transcripts were protein coding, representing 85.2%, 96.3%, and 90.6% of total altered transcripts in cytoplasm, nucleoplasm, and chromatin fractions, respectively. The second most-abundant major class of polyA RNAs following protein coding were lncRNAs (**Supplemental Figure S5d**).

We next asked whether the fractionated mRNA sequencing recapitulated the total transcriptome analysis (**Figure 1f**). We divided the total PRMT5i transcriptome analysis with PRMT5 inhibition into three categories: all significantly altered genes; upregulated; and downregulated. We then performed Fisher Exact tests comparing these gene sets to the significantly altered transcripts in the fractionated sequencing. After two-days of PRMT5 inhibition, the differentially expressed cytoplasmic transcripts significantly overlap with the total cell mRNA transcriptome (**Supplemental Figure S5e**). Furthermore, there was also a significant overlap with the total transcriptome up- and down-regulated transcripts. In contrast, for both the nucleoplasm- and chromatin-significantly altered transcripts, we observed that there was significant overlap only with the total and up-regulated genes from the total transcriptome experiment (**Supplemental Figure S5f-g**). These tests indicated that the total cell transcriptome mRNA-seq is predominantly measuring nuclear and chromatin RNA accumulation and may not represent productive transcription.

GO analysis of the chromatin enriched transcripts (p_adj_ < 0.05 and log_2_ Fold Change >= 0.58) identified RNA splicing, and ncRNA and rRNA processing ontologies, suggesting that PRMT5 regulates proper maturation and export of RNA processing transcripts (**Figure 6g**). GO analysis by cellular component ontology also highlighted transcripts involved in nuclear speckles and spliceosome complexes (**Supplemental Figure S5h**), further signifying PRMT5’s role in regulating nuclear substructures and RMA processing. Of note, these chromatin-enriched ontology analyses parallel that of the upregulated in the bulk RNA-seq and down regulated in nascent sequencing (**Figure 1h**).

To understand the normal compartmentalization of polyadenylated RNAs, we plotted the log_10_ spike-in normalized counts (TPM) of the cytoplasmic vs chromatin fractions (**Figure 6h**) and nucleoplasm vs chromatin fractions (**Supplemental Figure S5i**). Consistent with prior studies, we observed that certain subsets of RNAs have higher relative counts on chromatin whereas others predominantly are enriched in the cytoplasm.^58^ This indicated that at baseline, certain mature transcripts are more likely to associate with chromatin. We next subsetted the transcripts into the top and bottom 50% of TPM counts. We calculated the log_2_ of the chromatin to cytoplasm TPM ratio of our observed chromatin enriched transcripts upon PRMT5 inhibition vs. all expressed transcripts within the chromatin fraction (**Figure 6i and Supplemental Figure S5j**). We used a Kolmogorov-Smirnov (KS) test to compare the distribution and a Wilcoxon rank sum test of the lower expressing transcript ratios; the chromatin enriched transcripts had a significantly higher chromatin to cytoplasm ratio (p = 6.04×10^−25^ and 2.18×10^−27^, respectively). The higher expressing transcripts had a much less significant change in chromatin to cytoplasm ratio (p=5.6×10^−5^). We concluded that many polyadenylated transcripts are predisposed to interact with chromatin over cytoplasmic export.

### Mature transcript retention on chromatin is due to slower splicing and presence of retained introns

As we established that SNRPB chromatin accumulation was RNA-dependent, pICln knockdown recapitulated RNA accumulation on chromatin, and the well-defined role of PRMT5 in regulating splicing— particularly intron retention—we further examined the relationship between RNA splicing and mRNA chromatin accumulation. We observed that PRMT5-inhibition dependent chromatin-enriched transcripts were longer (p< 2.2×10^−16^) (**Supplemental Figure S6a**) and contained significantly more introns (p< 2.2×10^−16^) (**Figure 7a**) compared to the distributions of all expressed genes in A549 cells. To test whether a gene containing introns predicted its regulation by PRMT5 or being stuck on chromatin, we calculated an Odds Ratios of how each transcript was altered in these data sets. We included the Type I PRMT inhibited (MS023) total transcriptome as an additional control. Strikingly, we found that if a transcript has zero introns, it is unlikely to be either regulated by PRMT5 inhibition or trapped on chromatin. In contrast, if a transcript has one or more introns, it is more likely to be both regulated by and enriched on chromatin upon PRMT5 inhibition (**Figure 7b**).

**Figure 7.**
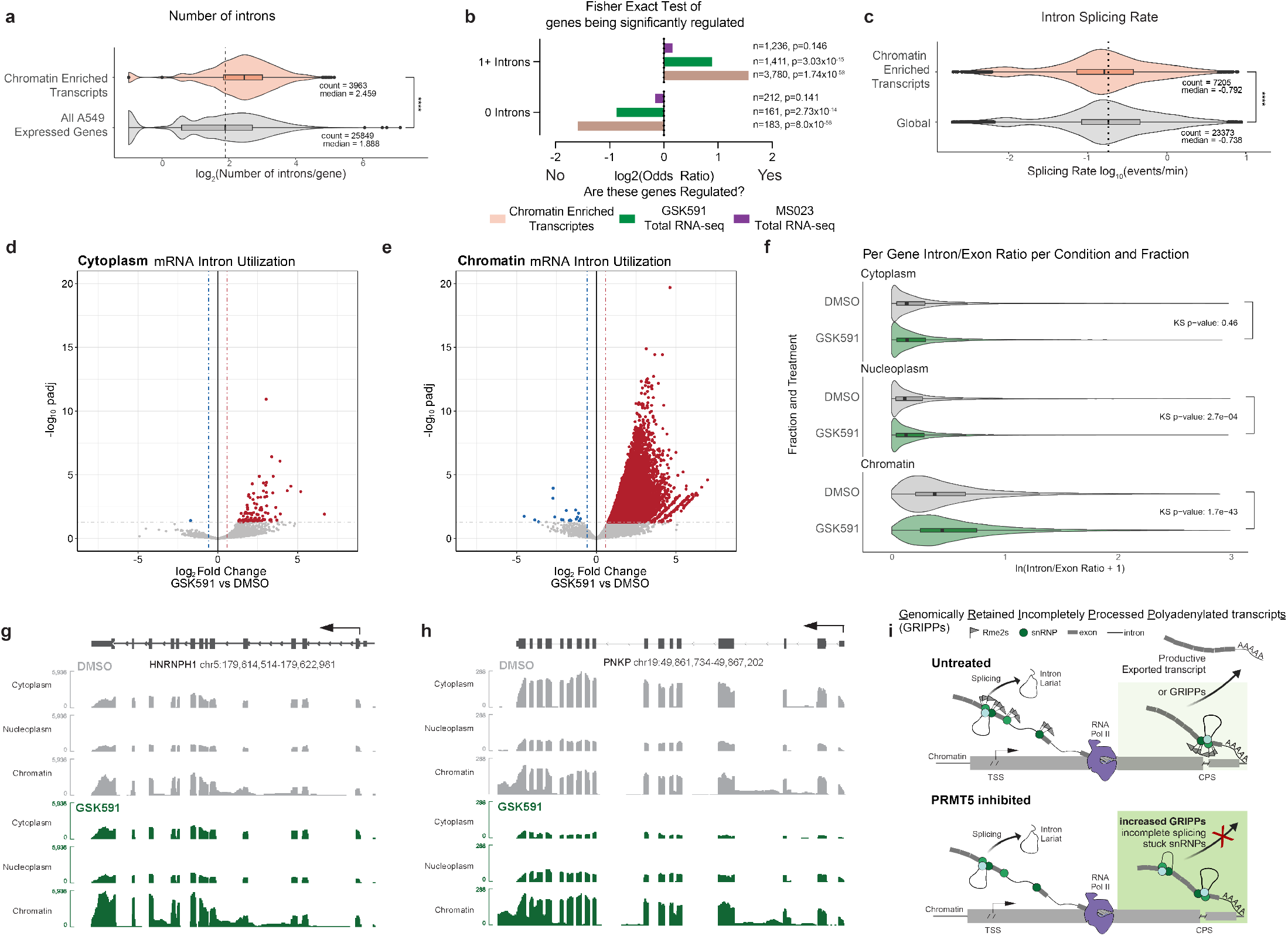
Chromatin enriched transcripts are defined by slower splicing rate and retained introns. **a)** Number of introns per gene for chromatin enriched transcripts and all expressed genes in A549 cells. Average compared by Wilcoxon Ranked Sum test. **** signifies p < 0.0001. **b)** Fisher Exact Test and ODDS ratio of chromatin enriched transcripts, PRMT5i altered transcripts, and Type I PRMTi altered transcripts depending on their number of introns. **c)** SKaTER-seq calculated splicing rate of chromatin enriched transcripts compared to global transcript splicing rates. Distribution compared with Kolmogorov-Smirnov test. **** signifies p < 0.0001. **d-e)** Volcano plots of intron utilization for introns in cytoplasm, and chromatin, compared to DMSO matched controls. Red introns have a log_2_FC>0.58 and a p_adj_>0.05. Blue introns have a log_2_FC<-0.58 and a p_adj_>0.05. **f)** Violin plots representing the average Intron/Exon ratio of genes significantly expressed in each cellular compartment. The distribution of ratios per compartment was tested using the Kolmogorov-Smirnov test. **g-h)** Selected transcript IGV tracks demonstrating scaled transcript and intron levels in cytoplasm, nucleoplasm, and chromatin for DMSO and GSK591 treated cells. **i)** Model figure illustrating Genomically Retained Incompletely Processed Polyadenylated Transcripts (GRIPPs) and their dependence on arginine methylation for productive escape from chromatin.

Notably, the Type I PRMT transcriptome is not significantly enriched in either scenario, indicating that Type I PRMTs have a separate, non-intron dependent regulatory mechanism of transcription.

To gain further understanding of the characteristics of the retained poly(A) RNAs, we analyzed data from our previous Splicing Kinetics and Transcript Elongation Rate (SKaTER)-seq study; in this work, we calculated rates of co-transcriptional processing, including RNA splicing, after PRMT5 inhibition.^8,59^ We observed no significant change in average global splicing rate upon PRMT5 inhibition (p=1) (**Supplemental Figure S6b**). Strikingly, when we compared global splicing rate to rates of our identified trapped mRNAs solved by the SKaTERseq algorithm, we observed that these transcripts are on average spliced significantly slower compared to the global population (p=1.44×10^−15^). Therefore, chromatin-enriched transcripts are both intron rich and slower to splice.

Next, we delineated how intron abundance varied between cellular compartments following PRMT5 inhibition. We calculated per intron read counts normalized to the exon counts within the same transcript for all significantly expressed genes (normalized counts > 5) from polyadenylated RNA. We then calculated differential intron abundance using the cross-species spike-in for normalization. In the cytoplasm, there were only 107 significantly retained introns for 94 unique genes (**Figure 7d**). These are true, “productive” retained introns as they have the potential to be translated. In the nucleoplasm, we identified 531 differentially observed introns from 440 genes (**Supplemental Figure S6c**). In the chromatin fraction there were 48,260 upregulated retained introns, corresponding to 10,219 unique genes (p_adj_<0.05, log_2_FoldChange>0.58) (**Figure 7e**). We found that the percentage of transcript introns throughout a transcript was higher in chromatin than in cytoplasmic or nucleoplasm transcripts (**Supplemental Figure S6d-f**). Similarly, the observed RI found on chromatin were evenly distributed throughout the transcripts (**Supplemental Figure S6g**). Furthermore, upon PRMT5 inhibition, we observe an increased distribution in the intron/exon ratio on chromatin upon PRM5 inhibition (p=1.7×10^−43^) (**Figure 7f**). PRMT5 inhibition had a modest increase in intronic reads/gene in the nucleoplasm (p=2.7×10^−4^) and had no significant difference in the cytoplasm (p=0.46). Despite the altered intron abundances (**Figure 7e**), the cytoplasmic intron:exon ration remained unchanged (**Figure 7f**). Taken together, we conclude that transcripts with retained introns are largely confined to the nucleus and are therefore detained.

Finally, we hypothesized that the PRMT5-induced, chromatin-enriched mRNAs would largely contain these increased RI reads. Strikingly, 81.5% of trapped transcripts have increased RI reads on chromatin. Looking specifically at genome tracks for chromatin-retained transcripts in PRMT5 inhibited cells compared to control, we observed two patterns: chromatin-retention with a corresponding depletion in nucleoplasm and cytoplasm, or chromatin-retention without a corresponding depletion in other cell compartments. This is exemplified by PNKP and HNRNPH1, respectively (**Figure 7g-h**). Overall, this demonstrates that chromatin-retained polyadenylated transcripts are rich in RIs. To better define the identified polyadenylated RNAs, we named this class of chromatin-enriched transcripts **G**enomically **R**etained **I**ncompletely **P**rocessed **P**olyadenylated Transcript**s** (GRIPPs). To test if chromatin-trapped transcripts had any impact on nascent transcription, we compared PRO-seq metagene profiles of GRIPPs and all A549 expressed transcripts (**Supplemental Figure S7h**). No gross differences were observed between these sets in promoter-proximal PRO-seq signal or in gene body signal. A cartoon model of our observations is shown in **Figure 7i**.

## Discussion

In this study we showed that through the methylation of SNRPB, PRMT5-pICln plays a pivotal role in regulating both mRNA splicing and chromatin escape. We demonstrate that PRMT5 inhibition leads to the significant retention of both mature polyadenylated mRNAs, lncRNAs, and snRNPs on chromatin, revealing a critical function of PRMT5 in post-transcriptional RNA processing. Our identification of GRIPPs—transcripts normally found enriched on chromatin and uncovered by PRMT5 inhibition—emphasizes the importance of PRMT5 in ensuring the efficient splicing and release of mRNAs from chromatin for subsequent nuclear export. Previous studies demonstrated that PRMT5 inhibition results in gross transcriptomic changes; however, few of these studies provide detailed mechanistic insight into the regulatory role of PRMT5.^1,2,11,60^ Our findings highlight the unique and non-redundant roles of different PRMTs in transcriptional regulation, with PRMT5 specifically influencing both nascent transcription and post-transcriptional processing.

As we observed significant transcriptional changes in total mRNA-seq after Type I PRMT and PRMT5 inhibition, our initial hypothesis was that histone modifications played an important role. The absence of any significant changes except for a modest increase in H3K27me3 after 7 days of PRMT5 inhibition was therefore quite surprising. Furthermore, there remains a lack of evidence that histone arginine methylation itself is abundant and critical for transcription in somatic tissues. While more conclusive studies are required, we were unable to observe any *bona fide* histone arginine methylation in our studies. We did observe one clear, confounding factor: histone acid extractions contain SNRP/Sm proteins, which themselves are heavily arginine methylated. Thus, SNRPB and SNRPD3 are likely the predominant bands in our immunoblots for Rme2s identified by previous studies from our group and likely from others.^1^

The concept of targeting PRMT5 adaptor proteins to modulate methylation of different PRMT5 substrates has been gaining traction as a possible therapeutic and also as an experimental test of PRMT5 function.^9,29^ As such, we decided to leverage knockdowns of these proteins to discern the mechanisms of both PRMT5 transcriptional regulation as well as our observed SNRPB chromatin accumulation. Consistent with prior studies, only pICln knockdown resulted in immunoblot changes consistent with SNRPB and SNRPD3 methylation. Similarly, only pICln knockdown replicated the splicing consequences of PRMT5 inhibition or knockdown. This highlighted the importance of Sm methylation in proper spliceosome function^.8,40^ Interestingly, RIOK1 knockdown had opposite effects on RI while affecting a shared subset of introns, consistent with reports that the pICln:RIOK1 ratio is critical in the regulation of proper mRNA splicing.^61^ Despite studies suggesting that CoPR5 enhances PRMT5 methylation of histones H3 and H4, we demonstrated that knockdown of CoPR5 resulted in minimal transcriptional changes^.9,13^

Both the total and fractionated proteomics analyses revealed that a major proteomic consequence of PRMT5 inhibition was the accumulation of SNRPB on chromatin. While there was a correlation between significantly altered protein and transcripts, there were many transcriptomic alterations that were not reflected in the proteomic analysis; this observation further suggested a potentially non-productive, post-transcriptional PRMT5 mechanism. We further demonstrated that SNRPB accumulated on chromatin upon loss of arginine methylation.

Surprisingly, but consistent with some prior studies, we did not find any clear evidence that loss of SNRPB methylation affected its assembly into mature snRNPs^.8,46^ Our findings suggest that the accumulation of SNRPB on chromatin upon PRMT5 inhibition is a direct consequence of the loss of arginine methylation. While the mechanism of enhanced SNRPB affinity for chromatin and RNA remains to be deciphered, we propose that modulation of RNA-protein interactions and cellular RNA homeostasis may be a fundamental role of arginine methylation.^62,63^

In keeping with this proposed mechanism, we demonstrated that PRMT5 inhibition resulted in RNA detention on chromatin. As dysregulation of chromatin-RNA interactions has significant implications in gene regulation and disease, including cancer ^17^, we further explored the relationship between RNA splicing and subcellular transport from transcription to nuclear export. The accumulation of GRIPPs on chromatin in the absence of PRMT5 activity points to a novel regulatory mechanism in which Sm symmetric dimethylarginine plays pivotal role in ensuring the timely processing and export of these transcripts. The significant enrichment of RNA processing and splicing-related transcripts among GRIPPs underscores a potential biological function of PRMT5 in maintaining RNA homeostasis and preventing genomic instability due to RNA-chromatin interactions. We also note that the use of a cross-species spike-in normalization control for the fractioned cell RNA-sequencing was essential for robust identification of true observations. Therefore, comparative studies conducted in the absence of spike-in control should be interpreted with appropriate skepticism.

To understand determinants of what makes a GRIPP transcript, we were struck by the strong connection between splicing and their presence on chromatin. This is exemplified by the correlation between the number of introns in a transcript and its likelihood of being detained on chromatin upon PRMT5 inhibition. GRIPPs, on average, had significant but modestly slower splicing rates than do all other transcripts.^8^ We conclude that GRIPPs consist of more complex transcripts, with a higher number of longer introns that are more susceptible to chromatin detention upon PRMT5 inhibition. A broad conclusion relevant across all PRMT5 mechanistic studies is that loss of Sm protein methylation and concomitant transcript trapping on chromatin may be in part responsible for observed total cell transcriptional phenotypes.

The identification of GRIPPs in our study suggests a previously unrecognized layer of RNA homeostasis that could play a significant role in normal cellular physiology. In the absence of PRMT5 inhibition, GRIPPs might function as a regulatory checkpoint within the nucleus, ensuring that only fully processed and correctly spliced mRNAs are exported to the cytoplasm for translation. This checkpoint mechanism could help maintain the fidelity of gene expression by preventing the premature release of incomplete or improperly spliced transcripts, which could otherwise lead to the production of aberrant proteins or non-functional RNA species. Additionally, GRIPPs might be involved in the fine-tuning of gene expression in response to cellular signals and stress. By temporarily retaining certain transcripts on chromatin, cells could rapidly adjust their gene expression profiles without the need for new transcription, allowing for a swift response to changing environmental conditions. This mechanism could be particularly important in processes requiring rapid adaptation, such as during cell differentiation, stress responses, and immune activation. Moreover, the association of GRIPPs with RNA processing and splicing-related transcripts suggests that they might play a role in the regulation of RNA processing machinery itself. By retaining these transcripts, cells might regulate the availability and activity of splicing factors and other RNA-binding proteins, thereby modulating the splicing landscape according to cellular requirements.

In summary, our study provides new understanding of the molecular mechanisms by which PRMT5 regulates RNA processing and chromatin interactions. Our findings highlight the essential role of PRMT5-mediated methylation in facilitating the proper splicing and release of mRNA from chromatin to maintain cellular RNA homeostasis. PRMT5-regulation of GRIPP transcript retention on chromatin represents a sophisticated mechanism by which cells ensure the integrity of their transcriptome and may rapidly respond to internal and external cues. Overall, our study expands the broader understanding of RNA-chromatin dynamics and emphasizes the importance of post-translational modifications in regulating gene expression and RNA metabolism.

### Limitations of This Study

While we robustly tested SNRPB accumulation on chromatin in a PRMT5 and methylarginine-dependent fashion, further studies will be needed to explore how unmethylated Sm proteins interact with GRIPPs and trap them on chromatin. Additionally, while we demonstrated that unmethylated SNRPB and SNRPD3 do assemble into mature snRNPs, this does not exclude the possibility that they form unique oligomeric complexes or have altered proteomic interactions upon loss of methylation. Further studies will explore how PRMT5-methylation affects SNRPB-protein interactions. Lastly, while we have characterized GRIPPs as a novel transcript class, further studies will explore their genomic characteristics and how they are protected from RNA degradation by the RNA exosome complex and prevented from RNA nuclear export.

## Methods

### Cell culture

A549 cells were purchased from ATCC for this study. Both A549 and HEK293T cells were cultured in DMEM (Corning), supplemented with 10% FBS (Hyclone), 100 I.U./mL penicillin and 100 µg/mL streptomycin. IMR90-hTERT were cultured in EMEM using the above additions. NSC-34 cells were cultured in DMEM, supplemented with 10% FBS and 1% 200 mM L-glutamine solution (Gibco). All cells were maintained at 37°C with 5% CO_2_. All cells were routinely tested for mycoplasma via PCR.^64^ Treatment with small molecule inhibitors dissolved in DMSO maintained final concentration of 0.01% DMSO, with 1 μM GSK591 (Cayman) or 1 μM MS023 (Cayman).

### Poly(A)-RNA sequencing

RNA was extracted using RNEasy Mini Kit (Qiagen) or Trizol (Invitrogen) for the cofactor knockdown and cellular fractionation experiments, respectively, as per the manufacturer’s protocols. RNA quantification and quality control were accomplished with the Qubit RNA BR Assay Kit (Invitrogen) and the Bioanalyzer 2100 (Agilent Technologies). Unstranded (PRMT inhibition time course) and stranded libraries (PRMT5 inhibition subcellular fractionation) were created by Novogene Genetics US. Barcoded libraries were sequenced on an Illumina platform using 150 nt paired-end libraries generating approximately 30 million paired-end reads (9GB total data) per biological replicate. Reads were trimmed and aligned to the human genome (hg38) or to a custom combined human and *Drosophila melanogaster* genome (hg38-dm6) using the nf-core/rnaseq pipeline.^65^ Differential abundance analysis for non-spike in studies was performed using DESeq2 with a custom script (https://github.com/Shechterlab/nextflow-rnaseq) and for the spike-in studies with a different script (https://github.com/Shechterlab/DESeq2_with_SpikeInNormalization). Alternative splicing events were determined using Replicate Multivariate Analysis of Transcript Splicing (rMATS, version 4.0.2).^66^

### Precision Run-On (PRO) nascent transcriptome sequencing

PRO-seq was performed as previously described.^67^

### Acid extraction of chromatin

This protocol was adapted from our previously published protocol.^27^ Hypotonic Lysis Buffer (HLB) was made by mixing 10 mM Tris-HCl pH 8.0, 1 mM KCl, 1.5 mM MgCl_2_, and 1 mM DTT, and supplemented with protease and phosphatase inhibitors. The Nuclear Lysis Buffer (NLB) was prepared with 10 mM Tris-HCl pH 8.0, 250 mM KCl, 0.1% NP40, and 1 mM DTT, and supplemented with protease and phosphatase inhibitors. Cell pellets were resuspended in HLB and incubated for 30 minutes to promote hypotonic swelling and mechanical lysis, followed by centrifugation to pellet intact nuclei. Nuclei were washed with additional HLB. Nuclei were then resuspended in NLB and rotated for 30 min at 4°C. Chromatin was then collected via centrifugation and washed with NLB.

Chromatin was then resuspended in 0.4N H_2_SO_4_, incubated for 30 min., then spun at 16,000xg for 10 min. at 4°C to remove nuclear debris. The supernatant, containing histones and acid-soluble proteins, was transferred to a fresh tube and treated with TCA to precipitate histones, which were then pelleted, washed with ice-cold acetone, and air-dried. The histone pellet was dissolved in an appropriate volume of ddH_2_O, characterized for concentration, and analyzed further using SDS/PAGE gel.

### Histone PTM mass spectrometry

Histones were acid-extracted from chromatin using the above protocol. Histone pellets were dissolved in 50 mM ammonium bicarbonate, pH 8.0, and histones were subjected to derivatization using 5 µL of propionic anhydride and 14 µL of ammonium hydroxide (all Sigma Aldrich) to balance the pH at 8.0. The mixture was incubated for 15 min and the procedure was repeated. Histones were then digested with 1 µg of sequencing grade trypsin (Promega) diluted in 50mM ammonium bicarbonate (1:20, enzyme:sample) overnight at room temperature. Derivatization reaction was repeated to derivatize peptide N-termini. The samples were dried in a vacuum centrifuge.

Prior to mass spectrometry analysis, samples were desalted using a 96-well plate filter (Orochem) packed with 1 mg of Oasis HLB C-18 resin (Waters). Briefly, the samples were resuspended in 100 µl of 0.1% TFA and loaded onto the HLB resin, which was previously equilibrated using 100 µl of the same buffer. After washing with 100 µl of 0.1% TFA, the samples were eluted with a buffer containing 70 µl of 60% acetonitrile and 0.1% TFA and then dried in a vacuum centrifuge.

Samples were resuspended in 10 µl of 0.1% TFA and loaded onto a Dionex RSLC Ultimate 300 (Thermo Scientific), coupled online with an Orbitrap Fusion Lumos (Thermo Scientific). Chromatographic separation was performed with a two-column system, consisting of a C-18 trap cartridge (300 µm ID, 5 mm length) and a picofrit analytical column (75 µm ID, 25 cm length) packed in-house with reversed-phase Repro-Sil Pur C18-AQ 3 µm resin. Peptides were separated using a 30 min gradient from 1-30% buffer B (buffer A: 0.1% formic acid, buffer B: 80% acetonitrile + 0.1% formic acid) at a flow rate of 300 nl/min. The mass spectrometer was set to acquire spectra in a data-independent acquisition (DIA) mode. Briefly, the full MS scan was set to 300-1100 m/z in the orbitrap with a resolution of 120,000 (at 200 m/z) and an AGC target of 5×10e5. MS/MS was performed in the orbitrap with sequential isolation windows of 50 m/z with an AGC target of 2×10e5 and an HCD collision energy of 30.

Histone peptides raw files were imported into EpiProfile 2.0 software.^68^ From the extracted ion chromatogram, the area under the curve was obtained and used to estimate the abundance of each peptide. To achieve the relative abundance of post-translational modifications (PTMs), the sum of all different modified forms of a histone peptide was considered as 100% and the area of the particular peptide was divided by the total area for that histone peptide in all of its modified forms. The relative ratio of two isobaric forms was estimated by averaging the ratio for each fragment ion with different mass between the two species. The resulting peptide lists generated by EpiProfile were exported to Microsoft Excel and further processed for a detailed analysis.

### Immunoblots

Whole cell lysis was performed using RIPA buffer (1% NP-40, 150 mM NaCl, 1mM EDTA, 50 mM Tris-HCl pH 8 at 4°C, 0.25% Sodium Deoxycholate, 0.1% SDS, protease inhibitor, and phosphatase inhibitor). Immunoblots were performed on PVDF (Immobilon, Millipore), detected by ECL chemiluminescence (Lumigen, TMA-6) and imaged with an ImageQuant LAS4000 (GE). A list of antibodies used is presented in **Supplemental Table S5**.

### Lentiviral CRISPRi knockdown

A two-plasmid lentiviral approach was used for knockdown of target genes. The lenti_dCas9-KRAB-MeCP2 plasmid was a gift from Andrea Califano (Addgene plasmid # 122205; http://n2t.net/addgene:122205; RRID:Addgene_122205) and pXPR_050 was a gift from John Doench & David Root (Addgene plasmid # 96925; http://n2t.net/addgene:96925; RRID:Addgene_96925).^69,70^ Custom sgRNA sequences (Supplementary file #) were designed using the Broad CRISPick software (https://portals.broadinstitute.org/gppx/crispick/public) and then cloned into pXPR_050 as previously described.^70^ Lentiviral particles were generated and transduced into A549 cells as previously done in our group.^8^ Stable expression of the dCas9-KRAB-MeCP2 construct was achieved in A549 cells with Blasticidin (Cayman) selection at 10 ug/mL before subsequent transduction with the sgRNA and selection with Puromycin at 2 ug/mL (Cayman). A list of sgRNAs used in this study are listed in **Supplemental Table S6**.

### Immunoprecipitation

After treatment, cells were harvested with trypsin (Corning), washed twice with 4°C PBS, and resuspended in modified RIPA buffer (0.5% NP-40, 150 mM NaCl, 1mM EDTA, 50 mM Tris-HCl pH 8 at 4°C) supplemented with 40 U/mL RNaseOUT (Thermo Fisher Scientific) and protease inhibitor (Thermo Fisher Scientific) at 10x cell pellet volume. Samples were incubated on ice for 10 min followed by sonication for 3×5 sec at 20% amplitude with a 1/8-inch probe-tip sonicator. All subsequent steps were completed at 4°C: Insoluble material was removed by 10,000g centrifugation for 10 min and lysates were pre-cleared by incubating with Protein G agarose (Millipore-Sigma, 16-201) for 30 minutes with rotation; supernatant was collected from agarose by centrifugation at 10,000g for 10 minutes. Concentrations of the cleared supernatants were normalized by bicinchoninic acid assay (Pierce) to 750 μg per condition (1.5 ug/uL). Primary antibody targeting Tri-methyl Guanosine (MBL Life Science, RN019M, clone 235-1; 5ug) was added and incubated over-night with rotation. Samples were treated with Protein G agarose and rotated for 2 hours, then washed three times by pelleting at 10,000g for 30 seconds and resuspending in fresh lysis buffer. Samples were prepared in 1x Laemmli buffer for western blotting with antibodies as specified.

### Northern Blotting

RNA was isolated using TRIzol (Thermo) and coprecipitated with GlycoBlue (Thermo). The RNA pellet was resuspended in sample buffer (6.8 M Urea in TBE with 10% glycerol and 0.25% Bromophenol Blue/Xylene Cyanide), heated at 90 °C for 3 minutes, followed by loading onto an 8% Urea Gel (National Diagnostics) that was pre-run for 45 minutes at 45 watts. The gel was run for one hour at 45 watts in 1x TBE (100 mM Tris, 100 mM Boric acid, and 0.2 mM EDTA) followed by transfer to nitrocellulose in 0.5x TBE at 30 mAmps for four hours at 4 °C. Following transfer, RNA was crosslinked to the membrane at 120,000 μJ/cm2 (UV Stratalinker 1800). 5’ end labeling of snRNA probes was performed using ^32^P-γ-ATP (PerkinElmer) with T4 PNK reaction (NEB). Unincorporated ^32^P-γ-ATP was removed using a Microspin G-25 column (Cytiva). Post-transfer hybridization was performed at 37 °C in a hybridization oven over-night with gentle agitation in hybridization buffer containing 100 mM NaHPO4 pH 7.2, 750 mM NaCl, 1x Denhardt’s Solution (0.02% BSA, 0.02% Ficoll 400, 0.02% Polyvinylpyrrolidone), 1% Herring sperm DNA, and 7% SDS. The next morning, the hybridization solution was carefully discarded according to institutional protocol and the membrane was washed twice with wash buffer (40 mM NaHPO4 pH 7.2, 2% SDS, 1 mM EDTA).

The wash buffer was also discarded according to institutional guidelines. The membrane was left to expose on a Phosphoimager screen (Cytiva) and imaged at 633 nm using a Typhoon 9400 Variable Mode Imager.

### Chromatin associated poly(A)-RNA protein enrichment and immunoblotting

Poly(A)-RNA isolation was performed with modifications to a previously described protocol ^71^. A549 cells were grown in the presence of 0.01% DMSO, 1 µM GSK591 (Cayman), or 1 µM MS023 (Cayman) for two days. Cells were washed with 4 °C PBS and irradiated on ice with 100 mJ cm-2 in a UV Stratalinker 1800. Cells were centrifuged at 500 x g for 10 minutes at 4 °C. Chromatin was isolated with nuclear lysis buffer (NLB; 10 mM Tris-HCl pH 7.5 at 4°C, 0.1% NP-40, 400 mM KCl, 1 mM DTT) supplemented with 40 U/mL RNaseOUT (Thermo), protease inhibitor (Thermo), and phosphatase inhibitor (Thermo). The chromatin pellet was resuspended in NLB and sonicated for five seconds at 20% amplitude with a probe-tip sonicator using a 1/8” tip. The sonicate was centrifuged at 10,000 x g for 10 minutes and the soluble material transferred to a low-adhesion RNase-free microcentrifuge tube. An aliquot from each sample was saved to serve as the unenriched control. The samples were split into two separate tubes, one of which received 10 µg of competitor 25-nt poly(A) RNA. Magnetic oligo-(dT) beads (NEB) were equilibrated in NLB and added to the enrichments. The samples were vortexed at room temperature for 10 minutes. The beads were then captured on a magnetic column, and the supernatant transferred to fresh tube for additional rounds of depletion. The beads were washed once with buffer A (10 mM Tris pH 7.5, 600 mM KCl, 1 mM EDTA, 0.1 % Triton X-100), followed by buffer B (10 mM Tris pH 7.5, 600 mM KCl, 1 mM EDTA) and lastly buffer C (10 mM Tris pH 7.5, 200 mM KCl, 1 mM EDTA). The RNA was eluted by incubating the beads in 10 µL of 10 mM Tris pH 7.5 at 80 °C for two minutes, capture of magnetic beads using a magnetic column, and quickly transferring the supernatant to a new tube. Protein from each condition and fraction was then immunoblotted with antibodies as indicated.

### Cellular Fractionation for Proteomics

This cellular fractionation was done as previously described.^8^ Briefly, cells were harvested via trypsinization post treatment and washed with PBS. All subsequent steps were performed at 4°C. Cells were suspended in 300 µL of Hypotonic Lysis Buffer (10 mM Tris-HCl pH 8 at 4°C, 0.1% NP-40, 1 mM KCl, 1.5 mM MgCl_2_, supplemented with protease inhibitor, phosphatase inhibitor, and 1µM of appropriate PRMT inhibitor) and rotated at for 30 min. Nuclei were spun down at 10,000xg for 10 min and the cytoplasm was collected. Nuclei were washed with Hypotonic Lysis Buffer. Nuclei were then resuspended in 300 µL of Nuclear Lysis Buffer (10 mM Tris-HCl pH 8 at 4°C, 0.1% NP-40, 400 mM KCl, 1 mM DTT, supplemented with protease inhibitor, phosphatase inhibitor, and 1µM of appropriate PRMT inhibitor) and rotated for 30 min. Chromatin was spun down at 10,000xg for 10 min and nucleoplasm was collected. The chromatin pellet was washed with additional Nuclear Lysis Buffer. Chromatin was resolubilized with prove-tip sonication at 20% amplitude for 5s. Total proteomic mass spectrometry was then performed on the cellular fractions.

### Total and fractionated proteomic mass spectrometry

Cells were homogenized with a probe sonicator in a buffer containing 5% SDS, 5 mM DTT and 50 mM ammonium bicarbonate (pH = 8), and left on the bench for about 1 hour for disulfide bond reduction. Samples were then alkylated with 20 mM iodoacetamide in the dark for 30 minutes. Afterward, phosphoric acid was added to the sample at a final concentration of 1.2%. Samples were diluted in six volumes of binding buffer (90% methanol and 10 mM ammonium bicarbonate, pH 8.0). After gentle mixing, the protein solution was loaded to an S-trap filter (Protifi) and spun at 500 g for 30 sec. The sample was washed twice with binding buffer. Finally, 1 µg of sequencing grade trypsin (Promega), diluted in 50 mM ammonium bicarbonate, was added into the S-trap filter and samples were digested at 37°C for 18 h. Peptides were eluted in three steps: (i) 40 µl of 50 mM ammonium bicarbonate, (ii) 40 µl of 0.1% TFA and (iii) 40 µl of 60% acetonitrile and 0.1% TFA. The peptide solution was pooled, spun at 1,000 g for 30 sec and dried in a vacuum centrifuge. Peptides were then desalted as performed for the histone peptides.

Samples were resuspended in 10 µl of 0.1% TFA and loaded onto a Dionex RSLC Ultimate 300 (Thermo Scientific), coupled online with an Orbitrap Fusion Lumos (Thermo Scientific). Chromatographic separation was performed with a two-column system, consisting of a C-18 trap cartridge (300 µm ID, 5 mm length) and a picofrit analytical column (75 µm ID, 25 cm length) packed in-house with reversed-phase Repro-Sil Pur C18-AQ 3 µm resin. Peptides were separated using a 90 min gradient from 4-30% buffer B (buffer A: 0.1% formic acid, buffer B: 80% acetonitrile + 0.1% formic acid) at a flow rate of 300 nl/min. The mass spectrometer was set to acquire spectra in a data-dependent acquisition (DDA) mode. Briefly, the full MS scan was set to 300-1200 m/z in the orbitrap with a resolution of 120,000 (at 200 m/z) and an AGC target of 5×10^5^. MS/MS was performed in the ion trap using the top speed mode (2 secs), an AGC target of 1×10^4^ and an HCD collision energy of 35.

Proteome raw files were searched using Proteome Discoverer software (v2.5, Thermo Scientific) using SEQUEST search engine and the SwissProt human database (updated March 2024). The search for total proteome included variable modification of N-terminal acetylation, and fixed modification of carbamidomethyl cysteine. Trypsin was specified as the digestive enzyme with up to 2 missed cleavages allowed. Mass tolerance was set to 10 pm for precursor ions and 0.2 Da for product ions. Peptide and protein false discovery rate was set to 1%. For all proteomic studies, DEP2 (Differential Enrichment analysis of Proteomics data 2)^72^ was used for data analysis with a maximum of one missing value and either GSimp or *k*-nearest number data imputation and False Discovery Rate control with Strimmer’s q-value. Heatmaps were generated from the log_2_ of the centered intensity measurements for the top differentially abundant proteins across all measurements. ClusterProfiler was used for protein ontology analysis.^73^

### Lentiviral inducible protein overexpression

SNRPB constructs were derived from plasmids created by Maron et al. and cloned into a Gateway Lentiviral system (Invitrogen).^8^ SmB WT, RtoA, and RtoK ORFs were individually PCR amplified and cloned into pDONR221 (Invitrogen) via a Gateway BP Clonase II Enzyme Mix (Invitrogen) using the manufacture’s protocol. Entry clones were then cloned into the destination vector, pLIX_403, via a Gateway LR Clonase II Enzyme Mix (Invitrogen). pLIX_403 was a gift from David Root (Addgene plasmid # 41395; http://n2t.net/addgene:41395; RRID:Addgene_41395). Lentiviral particles containing these constructs were generated and transduced into A549 cells as previously described.^8^ Transduced cells were selected with 2ug/mL Puromycin (Cayman) and maintained with 1ug/mL. Maximal protein induction was found after treating cells with Doxycycline at 1 ug/mL after three days (not shown).

### RNase treatment of Nuclei

Cells were pre-treated with 1 μM GSK591 for seven days. Cells were collected with trypsin, as described above. 5×10^6^ cells were lysed in 1 mL hypotonic lysis buffer (10 mM Tris pH 8 at 4°C, 1 mM KCl, 1.5 mM MgCl_2_, 0.1% NP-40, protease inhibitor cocktail, am 1 mM DTT) with end over end rotation for 30 min. Nuclei were pelleted at 10,000xg for 5 min. Nuclei were washed in 400 μL hypotonic lysis buffer and split in half. To one sample, 2 μg RNase A and 250 U RNase T1 were added; both samples were incubated at 37°C for 1 hour with intermitant mixing. Following incubation and at 4°C, nuclei were recovered by centrifugation at 10,000xg for 5 min. Nuclei were lysed with nuclear extraction buffer (see cellular fractionation for proteomics above) and chromatin was pelleted at 10,000 x g for 5 min. Nucleoplasm was collected as the supernatant. Chromatin was resuspended in 150 μL nuclear lysis buffer and solubilized with a tip sonicator. Nucleoplasm and chromatin fractions were then analyzed by immunoblotting.

### Subcellular RNA fractionation

This protocol followed previously described procedures with minor alterations.^8,19^,53,54 Briefly, following appropriate experimental treatments, cells were collected via a brief trypsinization followed by a PBS wash. Cells were then lysed in a hypotonic lysis buffer containing 10 mM Tris, 1 mM KCl, 1.5 mM MgCl2, 0.1% NP40, 0.5 mM DTT, and 0.5 mM EDTA, as well as 1x protease inhibitor and 40 U/ml RNaseOUT. Cells were rotated at 4°C for 15 minutes to promote lysis. The lysate was overlaid onto a 24% sucrose cushion, a density gradient solution consisting of 10 mM Tris, 15 mM KCl, 1 mM EDTA, 24% Sucrose (w/v), 0.5 mM DTT, 1x protease inhibitor, and 40 U/ml RNaseOUT, and centrifuged at 10,000xg for 10 min. Nuclei were washed in a PBS/1mM EDTA solution, and recollected by centrifuging at 10,000xg for 2 min. Nuclei were then resuspended in glycerol buffer, composed of 20 mM Tris, 75 mM NaCl, 0.5 mM EDTA, 50% Glycerol (v/v), 0.85 mM DTT, 40 U/ml RNaseOUT, and 1x protease inhibitor, and subsequently lysed using equal volume of nuclei lysis buffer, which contained 20 mM HEPES, 1 mM DTT, 7.5 mM MgCl2, 0.2 mM EDTA, 300 mM NaCl, 1% NP40, 1M Urea, 1x protease inhibitor, and 40 U/ml RNaseOUT. Following another PBS/EDTA wash, all three fractions were dissolved into TRIzol (Thermo Fisher Scientific), followed by subsequent isolation of RNA, DNA, and Protein as described in the manufacturer’s manual. RNA and DNA were subsequently quantified using the Qubit RNA BR Assay Kit and dsDNA BR Assay Kit, respectively (Invitrogen).

### Polyadenylated RNA quantification

Following Trizol purification, equal amounts (ng) of RNA were reverse transcribed using the Moloney Murine Leukemia Virus (MMLV) reverse transcriptase (Invitrogen) and oligo-dT primers. Residual RNA was removed via a RNase H digestion. cDNA was then quantified via the ssDNA Qubit Assay Kit (Invitrogen).

### Fractionated RNA-sequencing

The fractionation method used for the sequencing experiment is identical to the above subcellular RNA fractionation. Cells were counted in duplicate to obtain 5 million cells per condition and per biological replicate (collected on separate days). A batch of TRIzol was used with a spike in of total *D. melanogaster* S2 cells such that there are 250k (5% relative starting cell number) S2 cells per 750 uL of TRIzol. TRIzol was then aliquoted at 750 uL/tube and stored at -80°C until needed.

### Statistical analysis

All immunoblots were performed in at least two independent biological replicates. Statistical analysis was performed either using GraphPad Prism (v10.2.3) or R (v4.4.0). Wilcoxon Rank rank-sum test was used to compare the median between groups. Kolmogorov-Smirnov test was used to compare the distributions between groups. Independent t-tests were performed to compare means between only two groups. GeneOverlap^74^ with Fisher’s exact test was used to determine ORs of RI overlaps and transcript overlaps between different RNA-seq data sets.

## Supporting information

Table S1 - mRNASeq DEseq2 Output

Table S2 - PROSeq DEseq2 Output

Table S3 - Proteomic DEP2 Output

Table S4 - Fractionated mRNASeq DESeq2 Output

## Data Availability

Raw data for RNA-seq and PRO-seq is deposited under GEO (*upload pending*). Raw data for total cell and fractionated LC-MS/MS is deposited to the ProteomeXchange Consortium via the PRIDE partner repository (PXD054308). All code used to generate data in this manuscript can be found here: (https://github.com/Shechterlab/DeAngelo_2024).

## Acknowledgments

This work was supported by the National Institutes of Health [R01GM108646 to D.S., T32GM149364 to J.D.D., J.S.R., and A.M.S., and R01GM134379 to M.J.G.], Irma T. Hirschl and Monique Weill-Caulier Charitable Trusts to D.S., and The ALS Therapy Development Institute to D.S. The Sidoli lab gratefully acknowledges for funding the Hevolution Foundation (AFAR), the Einstein-Mount Sinai Diabetes center, and the NIH Office of the Director (S10OD030286). We thank Emmanuel Burgos for early studies.

## Author contributions

J.D.D.: conceptualization, investigation, analysis, visualization, and manuscript preparation; M.I.M.: conceptualization, investigation, analysis, visualization; J.S.R.: investigation and analysis; A.M.S.: investigation; V.G.: data analysis; S.S. and S.S. mass spectrometry investigation; J.B. and J.A. PRO-Seq investigation; M.J.G. conceptualization; D.S. conceptualization, analysis, visualization, manuscript preparation, and project supervision. To improve concision in portions of this manuscript, during the preparation of this work the authors used ChatGPT4.0o. After using this tool, the authors fully reviewed and edited the content as needed and take full responsibility for the content of the publication

## References

1. Maron, M.I., Lehman, S.M., Gayatri, S., DeAngelo, J.D., Hegde, S., Lorton, B.M., Sun, Y., Bai, D.L., Sidoli, S., Gupta, V., et al. (2021). Independent transcriptomic and proteomic regulation by type I and II protein arginine methyltransferases. iScience 24, 102971. 10.1016/j.isci.2021.102971.

2. Lorton, B.M., and Shechter, D. (2019). Cellular consequences of arginine methylation. Cell Mol Life Sci 76, 2933–2956. 10.1007/s00018-019-03140-2.

3. Guccione, E., and Richard, S. (2019). The regulation, functions and clinical relevance of arginine methylation. Nat Rev Mol Cell Biol 20, 642–657 10.1038/s41580-019-0155-x.

4. Friesen, W.J., Paushkin, S., Wyce, A., Massenet, S., Pesiridis, G.S., Van Duyne, G., Rappsilber, J., Mann, M., and Dreyfuss, G. (2001). The methylosome, a 20S complex containing JBP1 and pICln, produces dimethylarginine-modified Sm proteins. Mol Cell Biol 21, 8289–8300. 10.1128/MCB.21.24.8289-8300.2001.

5. Meister, G., Eggert, C., Buhler, D., Brahms, H., Kambach, C., and Fischer, U. (2001). Methylation of Sm proteins by a complex containing PRMT5 and the putative U snRNP assembly factor pICln. Curr Biol 11, 1990–1994. 10.1016/s0960-9822(01)00592-9.

6. Paknia, E., Chari, A., Stark, H., and Fischer, U. (2016). The Ribosome Cooperates with the Assembly Chaperone pICln to Initiate Formation of snRNPs. Cell Rep 16, 3103–3112. 10.1016/j.celrep.2016.08.047.

7. Urlaub, H., Raker, V.A., Kostka, S., and Luhrmann, R. (2001). Sm protein-Sm site RNA interactions within the inner ring of the spliceosomal snRNP core structure. EMBO J 20, 187–196. 10.1093/emboj/20.1.187.

8. Maron, M.I., Casill, A.D., Gupta, V., Roth, J.S., Sidoli, S., Query, C.C., Gamble, M.J., and Shechter, D. (2022). Type I and II PRMTs inversely regulate post-transcriptional intron detention through Sm and CHTOP methylation. Elife 11. 10.7554/eLife.72867.

9. Mulvaney, K.M., Blomquist, C., Acharya, N., Li, R., O’Keefe, M., Ranaghan, M., Stokes, M., Nelson, A.J., Jain, S.S., Columbus, J., et al. (2020). Molecular basis for substrate recruitment to the PRMT5 methylosome. bioRxiv, 2020.2008.2022.256347. 10.1101/2020.08.22.256347.

10. Pesiridis, G.S., Diamond, E., and Van Duyne, G.D. (2009). Role of pICLn in methylation of Sm proteins by PRMT5. J Biol Chem 284, 21347–21359. 10.1074/jbc.M109.015578.

11. Chen, H., Lorton, B., Gupta, V., and Shechter, D. (2017). A TGFbeta-PRMT5-MEP50 axis regulates cancer cell invasion through histone H3 and H4 arginine methylation coupled transcriptional activation and repression. Oncogene 36, 373–386. 10.1038/onc.2016.205.

12. Brahms, H., Meheus, L., de Brabandere, V., Fischer, U., and Luhrmann, R. (2001). Symmetrical dimethylation of arginine residues in spliceosomal Sm protein B/B’ and the Sm-like protein LSm4, and their interaction with the SMN protein. RNA 7, 1531–1542. 10.1017/s135583820101442x.

13. Lacroix, M., El Messaoudi, S., Rodier, G., Le Cam, A., Sardet, C., and Fabbrizio, E. (2008). The histone-binding protein COPR5 is required for nuclear functions of the protein arginine methyltransferase PRMT5. EMBO Rep 9, 452–458. 10.1038/embor.2008.45.

14. Guderian, G., Peter, C., Wiesner, J., Sickmann, A., Schulze-Osthoff, K., Fischer, U., and Grimmler, M. (2011). RioK1, a new interactor of protein arginine methyltransferase 5 (PRMT5), competes with pICln for binding and modulates PRMT5 complex composition and substrate specificity. J Biol Chem 286, 1976–1986. 10.1074/jbc.M110.148486.

15. Ren, J., Wang, Y., Liang, Y., Zhang, Y., Bao, S., and Xu, Z. (2010). Methylation of ribosomal protein S10 by protein-arginine methyltransferase 5 regulates ribosome biogenesis. J Biol Chem 285, 12695–12705. 10.1074/jbc.M110.103911.

16. Statello, L., Guo, C.J., Chen, L.L., and Huarte, M. (2021). Gene regulation by long non-coding RNAs and its biological functions. Nat Rev Mol Cell Biol 22, 96–118. 10.1038/s41580-020-00315-9.

17. Tang, J., Wang, X., Xiao, D., Liu, S., and Tao, Y. (2023). The chromatin-associated RNAs in gene regulation and cancer. Mol Cancer 22, 27. 10.1186/s12943-023-01724-y.

18. Li, X., and Fu, X.D. (2019). Chromatin-associated RNAs as facilitators of functional genomic interactions. Nat Rev Genet 20, 503–519. 10.1038/s41576-019-0135-1.

19. Yeom, K.H., Pan, Z., Lin, C.H., Lim, H.Y., Xiao, W., Xing, Y., and Black, D.L. (2021). Tracking pre-mRNA maturation across subcellular compartments identifies developmental gene regulation through intron retention and nuclear anchoring. Genome Res 31, 1106–1119. 10.1101/gr.273904.120.

20. Sollier, J., and Cimprich, K.A. (2015). Breaking bad: R-loops and genome integrity. Trends Cell Biol 25, 514–522. 10.1016/j.tcb.2015.05.003.

21. Boutz, P.L., Bhutkar, A., and Sharp, P.A. (2015). Detained introns are a novel, widespread class of post-transcriptionally spliced introns. Genes Dev 29, 63–80. 10.1101/gad.247361.114.

22. Gary, J.D., and Clarke, S. (1998). RNA and protein interactions modulated by protein arginine methylation. Prog Nucleic Acid Res Mol Biol 61, 65–131. 10.1016/s0079-6603(08)60825-9.

23. Duncan, K.W., Rioux, N., Boriack-Sjodin, P.A., Munchhof, M.J., Reiter, L.A., Majer, C.R., Jin, L., Johnston, L.D., Chan-Penebre, E., Kuplast, K.G., et al. (2016). Structure and Property Guided Design in the Identification of PRMT5 Tool Compound EPZ015666. ACS Med Chem Lett 7, 162–166. 10.1021/acsmedchemlett.5b00380.

24. Eram, M.S., Shen, Y., Szewczyk, M., Wu, H., Senisterra, G., Li, F., Butler, K.V., Kaniskan, H.U., Speed, B.A., Dela Sena, C., et al. (2016). A Potent, Selective, and Cell-Active Inhibitor of Human Type I Protein Arginine Methyltransferases. ACS Chem Biol 11, 772–781. 10.1021/acschembio.5b00839.

25. Liu, F., Xu, Y., Lu, X., Hamard, P.J., Karl, D.L., Man, N., Mookhtiar, A.K., Martinez, C., Lossos, I.S., Sun, J., and Nimer, S.D. (2020). PRMT5-mediated histone arginine methylation antagonizes transcriptional repression by polycomb complex PRC2. Nucleic Acids Res 48, 2956–2968. 10.1093/nar/gkaa065.

26. Kwak, H., Fuda, N.J., Core, L.J., and Lis, J.T. (2013). Precise maps of RNA polymerase reveal how promoters direct initiation and pausing. Science 339, 950–953. 10.1126/science.1229386.

27. Shechter, D., Dormann, H.L., Allis, C.D., and Hake, S.B. (2007). Extraction, purification and analysis of histones. Nat Protoc 2, 1445–1457. 10.1038/nprot.2007.202.

28. Mulvaney, K.M., Blomquist, C., Acharya, N., Li, R., Ranaghan, M.J., O’Keefe, M., Rodriguez, D.J., Young, M.J., Kesar, D., Pal, D., et al. (2021). Molecular basis for substrate recruitment to the PRMT5 methylosome. Mol Cell 81, 3481–3495 e3487. 10.1016/j.molcel.2021.07.019.

29. McKinney, D.C., McMillan, B.J., Ranaghan, M.J., Moroco, J.A., Brousseau, M., Mullin-Bernstein, Z., O’Keefe, M., McCarren, P., Mesleh, M.F., Mulvaney, K.M., et al. (2021). Discovery of a First-in-Class Inhibitor of the PRMT5-Substrate Adaptor Interaction. J Med Chem 64, 11148–11168. 10.1021/acs.jmedchem.1c00507.

30. Stopa, N., Krebs, J.E., and Shechter, D. (2015). The PRMT5 arginine methyltransferase: many roles in development, cancer and beyond. Cell Mol Life Sci 72, 2041–2059. 10.1007/s00018-015-1847-9.

31. Krzyzanowski, A., Gasper, R., Adihou, H., t Hart, P., and Waldmann, H. (2021). Biochemical Investigation of the Interaction of pICln, RioK1 and COPR5 with the PRMT5-MEP50 Complex. Chembiochem. 10.1002/cbic.202100079.

32. Yeo, N.C., Chavez, A., Lance-Byrne, A., Chan, Y., Menn, D., Milanova, D., Kuo, C.-C., Guo, X., Sharma, S., Tung, A., et al. (2018). An enhanced CRISPR repressor for targeted mammalian gene regulation. Nat Methods 15, 611–616. 10.1038/s41592-018-0048-5.

33. Ho, M.C., Wilczek, C., Bonanno, J.B., Xing, L., Seznec, J., Matsui, T., Carter, L.G., Onikubo, T., Kumar, P.R., Chan, M.K., et al. (2013). Structure of the arginine methyltransferase PRMT5-MEP50 reveals a mechanism for substrate specificity. PLoS One 8, e57008. 10.1371/journal.pone.0057008.

34. Wilczek, C., Chitta, R., Woo, E., Shabanowitz, J., Chait, B.T., Hunt, D.F., and Shechter, D. (2011). Protein arginine methyltransferase Prmt5-Mep50 methylates histones H2A and H4 and the histone chaperone nucleoplasmin in Xenopus laevis eggs. J Biol Chem 286, 42221–42231. 10.1074/jbc.M111.303677.

35. Lin, C.C., Chang, T.C., Wang, Y., Guo, L., Gao, Y., Bikorimana, E., Lemoff, A., Fang, Y.V., Zhang, H., Zhang, Y., et al. (2024). PRMT5 is an actionable therapeutic target in CDK4/6 inhibitor-resistant ER+/RB-deficient breast cancer. Nat Commun 15, 2287. 10.1038/s41467-024-46495-2.

36. Scoumanne, A., Zhang, J., and Chen, X. (2009). PRMT5 is required for cell-cycle progression and p53 tumor suppressor function. Nucleic Acids Res 37, 4965–4976. 10.1093/nar/gkp516.

37. Prusty, A.B., Meduri, R., Prusty, B.K., Vanselow, J., Schlosser, A., and Fischer, U. (2017). Impaired spliceosomal UsnRNP assembly leads to Sm mRNA down-regulation and Sm protein degradation. J Cell Biol 216, 2391–2407. 10.1083/jcb.201611108.

38. Fong, J.Y., Pignata, L., Goy, P.A., Kawabata, K.C., Lee, S.C., Koh, C.M., Musiani, D., Massignani, E., Kotini, A.G., Penson, A., et al. (2019). Therapeutic Targeting of RNA Splicing Catalysis through Inhibition of Protein Arginine Methylation. Cancer Cell 36, 194–209 e199. 10.1016/j.ccell.2019.07.003.

39. Sachamitr, P., Ho, J.C., Ciamponi, F.E., Ba-Alawi, W., Coutinho, F.J., Guilhamon, P., Kushida, M.M., Cavalli, F.M.G., Lee, L., Rastegar, N., et al. (2021). PRMT5 inhibition disrupts splicing and stemness in glioblastoma. Nat Commun 12, 979. 10.1038/s41467-021-21204-5.

40. Mateos, J.L., Sanchez, S.E., Legris, M., Esteve-Bruna, D., Torchio, J.C., Petrillo, E., Goretti, D., Blanco-Tourinan, N., Seymour, D.K., Schmid, M., et al. (2023). PICLN modulates alternative splicing and light/temperature responses in plants. Plant Physiol 191, 1036–1051. 10.1093/plphys/kiac527.

41. Jobert, L., Argentini, M., and Tora, L. (2009). PRMT1 mediated methylation of TAF15 is required for its positive gene regulatory function. Exp Cell Res 315, 1273–1286. 10.1016/j.yexcr.2008.12.008.

42. Matera, A.G., Terns, R.M., and Terns, M.P. (2007). Non-coding RNAs: lessons from the small nuclear and small nucleolar RNAs. Nat Rev Mol Cell Biol 8, 209–220. 10.1038/nrm2124.

43. Andersen, J., and Zieve, G.W. (1991). Assembly and intracellular transport of snRNP particles. Bioessays 13, 57–64. 10.1002/bies.950130203.

44. Pomeranz Krummel, D.A., Oubridge, C., Leung, A.K., Li, J., and Nagai, K. (2009). Crystal structure of human spliceosomal U1 snRNP at 5.5 A resolution. Nature 458, 475–480. 10.1038/nature07851.

45. Will, C.L., and Luhrmann, R. (2011). Spliceosome structure and function. Cold Spring Harbor perspectives in biology 3. 10.1101/cshperspect.a003707.

46. Gonsalvez, G.B., Praveen, K., Hicks, A.J., Tian, L., and Matera, A.G. (2008). Sm protein methylation is dispensable for snRNP assembly in Drosophila melanogaster. RNA 14, 878–887. 10.1261/rna.940708.

47. Boisvert, F.M., Cote, J., Boulanger, M.C., Cleroux, P., Bachand, F., Autexier, C., and Richard, S. (2002). Symmetrical dimethylarginine methylation is required for the localization of SMN in Cajal bodies and pre-mRNA splicing. J Cell Biol 159, 957–969. 10.1083/jcb.200207028.

48. Stanek, D. (2023). Coilin and Cajal bodies. Nucleus 14, 2256036. 10.1080/19491034.2023.2256036.

49. Roithova, A., Klimesova, K., Panek, J., Will, C.L., Luhrmann, R., Stanek, D., and Girard, C. (2018). The Sm-core mediates the retention of partially-assembled spliceosomal snRNPs in Cajal bodies until their full maturation. Nucleic Acids Res 46, 3774–3790. 10.1093/nar/gky070.

50. Hensold, J.O., Barth, D., and Stratton, C.A. (1996). RNA polymerase II inhibitor, 5,6-dichloro-1-beta-D-ribofuranosylbenzimidazole (DRB) causes erythroleukemic differentiation and transcriptional activation of erythroid genes. J Cell Physiol 168, 105–113. 10.1002/(SICI)1097-4652(199607)168:1<105::AID-JCP13>3.0.CO;2-6.

51. Rio, D.C., Ares, M., Jr., Hannon, G.J., and Nilsen, T.W. (2010). Purification of RNA using TRIzol (TRI reagent). Cold Spring Harb Protoc 2010, pdb prot5439. 10.1101/pdb.prot5439.

52. Khurana, A., Chadha, Y., and Schmoller, K.M. (2023). Too big not to fail: Different paths lead to senescence of enlarged cells. Mol Cell 83, 3946–3947. 10.1016/j.molcel.2023.10.024.

53. Pandya-Jones, A., and Black, D.L. (2009). Co-transcriptional splicing of constitutive and alternative exons. RNA 15, 1896–1908. 10.1261/rna.1714509.

54. Yeom, K.H., and Damianov, A. (2017). Methods for Extraction of RNA, Proteins, or Protein Complexes from Subcellular Compartments of Eukaryotic Cells. Methods Mol Biol 1648, 155–167. 10.1007/978-1-4939-7204-3_12.

55. Jiang, L., Schlesinger, F., Davis, C.A., Zhang, Y., Li, R., Salit, M., Gingeras, T.R., and Oliver, B. (2011). Synthetic spike-in standards for RNA-seq experiments. Genome Res 21, 1543–1551. 10.1101/gr.121095.111.

56. Schertzer, M.D., Murvin, M.M., and Calabrese, J.M. (2020). Using RNA Sequencing and Spike-in RNAs to Measure Intracellular Abundance of lncRNAs and mRNAs. Bio Protoc 10. 10.21769/bioprotoc.3772.

57. Taruttis, F., Feist, M., Schwarzfischer, P., Gronwald, W., Kube, D., Spang, R., and Engelmann, J.C. (2017). External calibration with Drosophila whole-cell spike-ins delivers absolute mRNA fold changes from human RNA-Seq and qPCR data. Biotechniques 62, 53–61. 10.2144/000114514.

58. Werner, M.S., Sullivan, M.A., Shah, R.N., Nadadur, R.D., Grzybowski, A.T., Galat, V., Moskowitz, I.P., and Ruthenburg, A.J. (2017). Chromatin-enriched lncRNAs can act as cell-type specific activators of proximal gene transcription. Nat Struct Mol Biol 24, 596–603. 10.1038/nsmb.3424.

59. Casill, A.D., Haimowitz, A.J., Kosmyna, B., Query, C.C., Ye, K., and Gamble, M.J. (2021). Spatial organization of transcript elongation and splicing kinetics. bioRxiv, 2021.2001.2028.428713. 10.1101/2021.01.28.428713.

60. Litzler, L.C., Zahn, A., Meli, A.P., Hebert, S., Patenaude, A.M., Methot, S.P., Sprumont, A., Bois, T., Kitamura, D., Costantino, S., et al. (2019). PRMT5 is essential for B cell development and germinal center dynamics. Nat Commun 10, 22. 10.1038/s41467-018-07884-6.

61. Braun, C.J., Stanciu, M., Boutz, P.L., Patterson, J.C., Calligaris, D., Higuchi, F., Neupane, R., Fenoglio, S., Cahill, D.P., Wakimoto, H., et al. (2017). Coordinated Splicing of Regulatory Detained Introns within Oncogenic Transcripts Creates an Exploitable Vulnerability in Malignant Glioma. Cancer Cell 32, 411–426 e411. 10.1016/j.ccell.2017.08.018.

62. Avila-Lopez, P., and Lauberth, S.M. (2024). Exploring new roles for RNA-binding proteins in epigenetic and gene regulation. Curr Opin Genet Dev 84, 102136. 10.1016/j.gde.2023.102136.

63. Vijayakumar, A., Majumder, M., Yin, S., Brobbey, C., Karam, J., Howley, B., Howe, P.H., Berto, S., Madan, L.K., Gan, W., and Palanisamy, V. (2024). PRMT5-mediated arginine methylation of FXR1 is essential for RNA binding in cancer cells. Nucleic Acids Res. 10.1093/nar/gkae319.

64. Dussurget, O., and Roulland-Dussoix, D. (1994). Rapid, sensitive PCR-based detection of mycoplasmas in simulated samples of animal sera. Appl Environ Microbiol 60, 953–959. 10.1128/aem.60.3.953-959.1994.

65. Ewels, P.A., Peltzer, A., Fillinger, S., Patel, H., Alneberg, J., Wilm, A., Garcia, M.U., Di Tommaso, P., and Nahnsen, S. (2020). The nf-core framework for community-curated bioinformatics pipelines. Nat Biotechnol 38, 276–278. 10.1038/s41587-020-0439-x.

66. Shen, S., Park, J.W., Lu, Z.X., Lin, L., Henry, M.D., Wu, Y.N., Zhou, Q., and Xing, Y. (2014). rMATS: robust and flexible detection of differential alternative splicing from replicate RNA-Seq data. Proc Natl Acad Sci U S A 111, E5593–5601. 10.1073/pnas.1419161111.

67. Mahat, D.B., Kwak, H., Booth, G.T., Jonkers, I.H., Danko, C.G., Patel, R.K., Waters, C.T., Munson, K., Core, L.J., and Lis, J.T. (2016). Base-pair-resolution genome-wide mapping of active RNA polymerases using precision nuclear run-on (PRO-seq). Nat Protoc 11, 1455–1476. 10.1038/nprot.2016.086.

68. Yuan, Z.F., Sidoli, S., Marchione, D.M., Simithy, J., Janssen, K.A., Szurgot, M.R., and Garcia, B.A. (2018). EpiProfile 2.0: A Computational Platform for Processing Epi-Proteomics Mass Spectrometry Data. J Proteome Res 17, 2533–2541. 10.1021/acs.jproteome.8b00133.

69. Tan, X., Worley, J., Turunen, M., Wong, K., Fernández, E.C., Paull, E., Jones, S., Wang, J., Noh, H., Salvatori, B., et al. (2022). Interrogation of genome-wide, experimentally dissected gene regulatory networks reveals mechanisms underlying dynamic cellular state control. bioRxiv, 2021.2006.2028.449297. 10.1101/2021.06.28.449297.

70. Sanson, K.R., Hanna, R.E., Hegde, M., Donovan, K.F., Strand, C., Sullender, M.E., Vaimberg, E.W., Goodale, A., Root, D.E., Piccioni, F., and Doench, J.G. (2018). Optimized libraries for CRISPR-Cas9 genetic screens with multiple modalities. Nat Commun 9, 5416. 10.1038/s41467-018-07901-8.

71. Iadevaia, V., Matia-González, A.M., and Gerber, A.P. (2018). An Oligonucleotide-based Tandem RNA Isolation Procedure to Recover Eukaryotic mRNA-Protein Complexes. Journal of Visualized Experiments. 10.3791/58223.

72. Zhang, X., Smits, A.H., van Tilburg, G.B., Ovaa, H., Huber, W., and Vermeulen, M. (2018). Proteome-wide identification of ubiquitin interactions using UbIA-MS. Nat Protoc 13, 530–550. 10.1038/nprot.2017.147.

73. Yu, G., Wang, L.G., Han, Y., and He, Q.Y. (2012). clusterProfiler: an R package for comparing biological themes among gene clusters. OMICS 16, 284–287. 10.1089/omi.2011.0118.

74. Shen, L. (2020). GeneOverlap: Test and visualize gene overlaps. https://github.com/shenlab-sinai/GeneOverlap.

