## Supplemental Figures S1-S6, Supplemental Tables S5, S6 for "Productive mRNA Chromatin Escape is Promoted by PRMT5 Methylation of SNRPB"

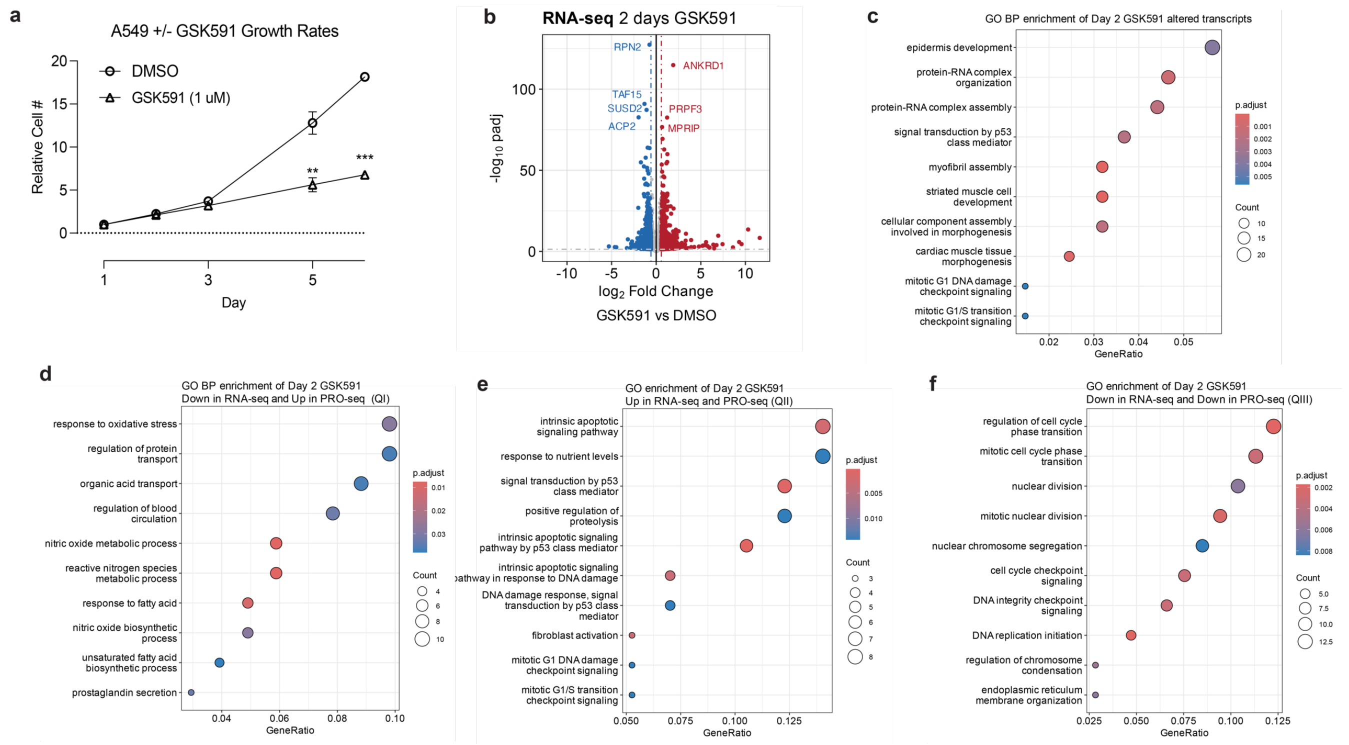


**Supplemental Figure S1 (Related to Figure 1). Additional PRMT5 inhibition characterization. a)** Growth curve of GSK591 treatment on A549 cells; **<0.01, ***<0.0001. **b)** Volcano plot of poly(A) RNA-sequencing following two days of GSK591 treatment, compared to DMSO control. **c)** Dot plot of GO Biological Processes (BP) terms for total mRNA sequencing of GSK591 treatment. **d-f)** Dot plots of GO Biological Processes (BP) terms for quadrants of linear correlation plot between GSK591 mRNA-sequencing and PRO-sequencing.

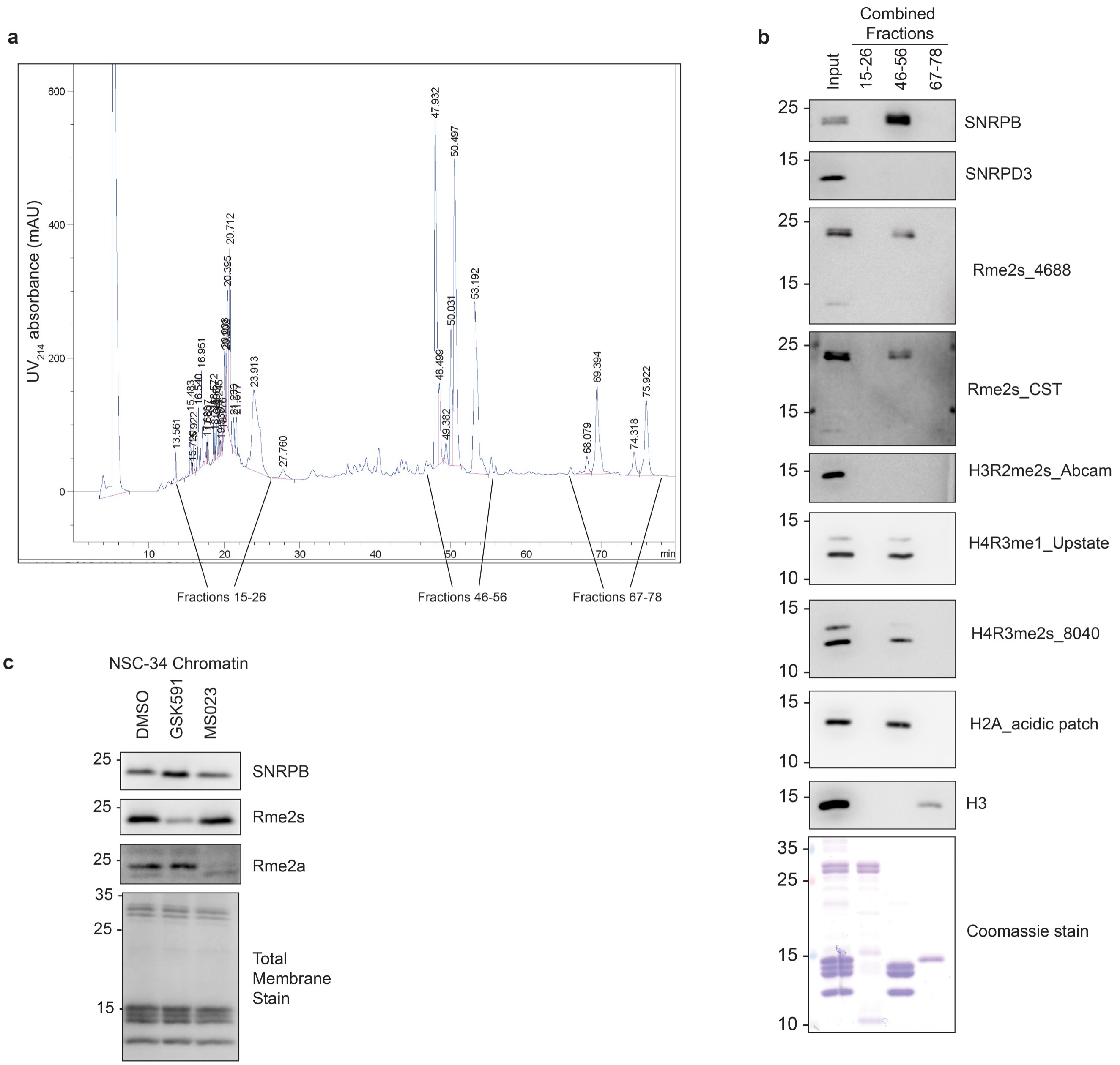


**Supplemental Figure S1 (Related to Figure 2). Sm proteins are a contaminant of acid extracted histones. a)** HPLC trace showing UV absorbance at 214nm for a run consisting of acid extracted chromatin. **b)** Western blot of combined fractions, analyzing presence of Sm proteins, histones, and selected histone PTMs. **c)** Immunoblot of chromatin fraction of NSC-34 cells following 2 day treatment of GSK591 or MS023.

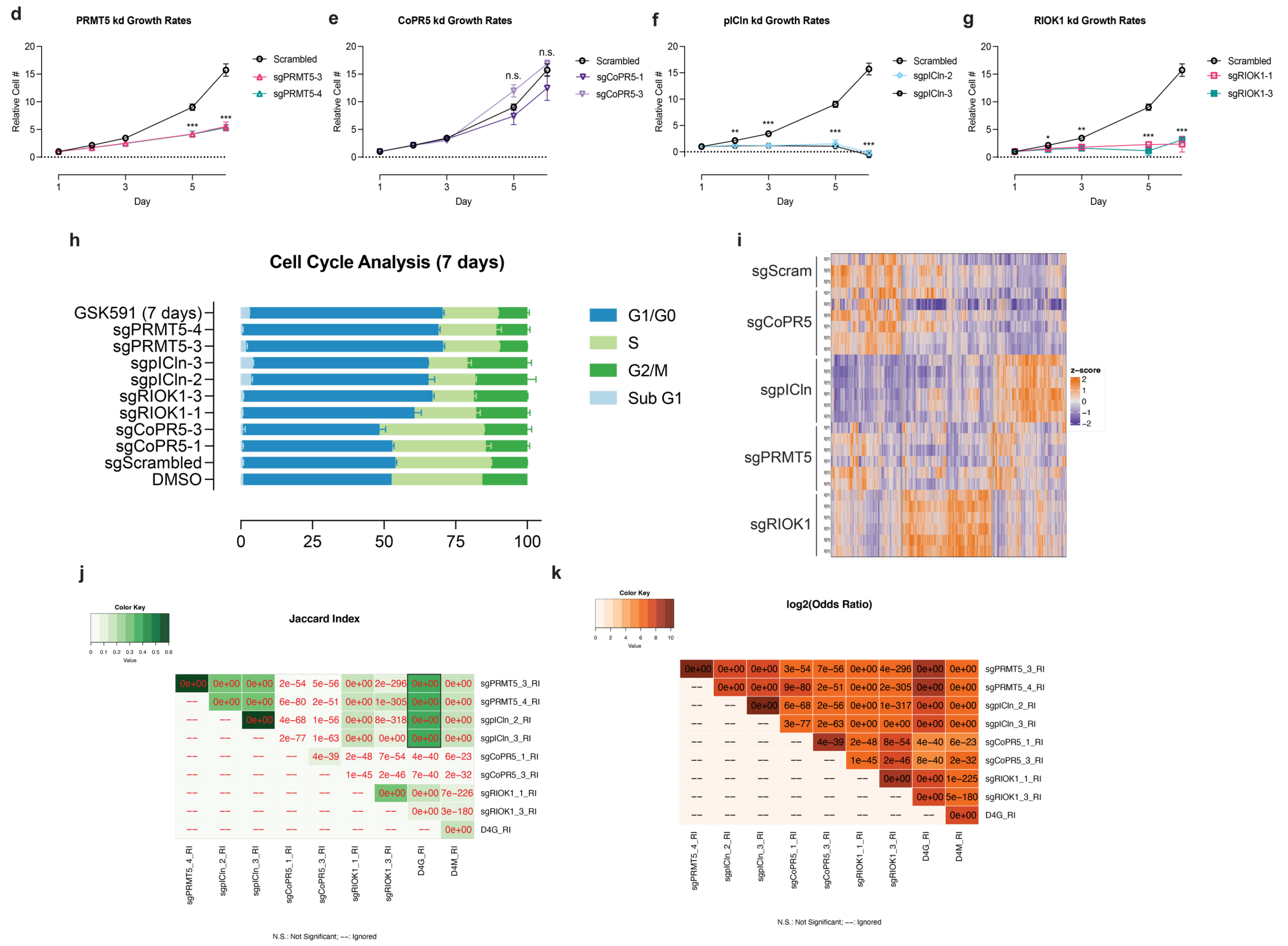


**Supplemental Figure S2 continued (Related to Figure 2). Characterization of PRMT5 and adaptor protein knockdown. d-g)** Growth curve of knockdown cell lines; *<0.01, **<0.01, ***<0.001, n.s. = not significant **h)** Cell Cycle analysis of PRMT5 and adaptor knockdowns using propidium iodide (PI) staining cell cytometry. **i)** Z-score heatmap TPM counts from RNA sequencing of PRMT5 and adaptor knockdowns. **j)** Heatmap showing the Jaccard Index for the overlap of differentially retained introns for PRMT5 and adaptor protein knockdowns. The p-value of each overlap is shown in the rectangles. Color is proportional to the percentage of intron overlap (Jaccard Index). **k)** Heatmap showing the Odds Ratio for the overlap of the differentially retained introns for PRMT5 and adaptor protein knockdowns. The p-value of the overlap is shows in each rectangle. Rectangle color is proportional to $log_2$(Odds Ratio).

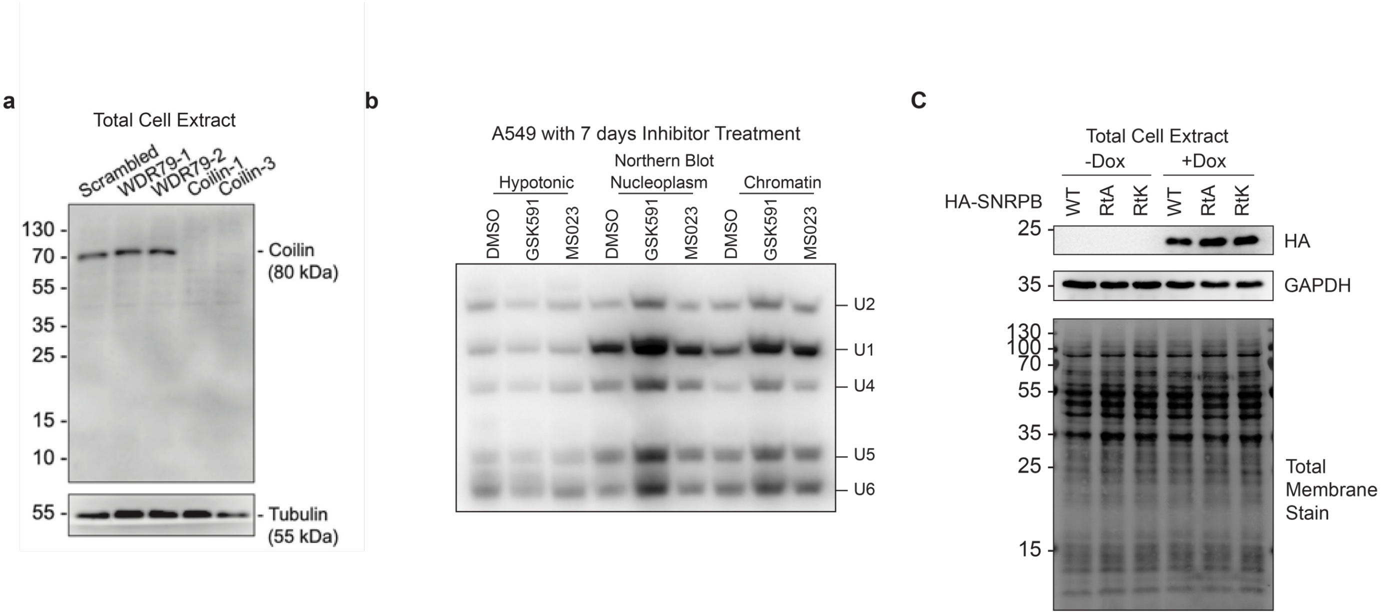


**Supplemental Figure S3 (Related to Figure 4). a)** Immunoblot controls of Coilin knockdown using CRISPRi in A549 cell. **b)** Northern blot testing snRNP abundance across cellular compartments after PRMT inhibition. **c)** Immunoblot of total cell lysate testing induction levels of mutant SNRPB proteins following four days of DOX (1ug/mL) induction.

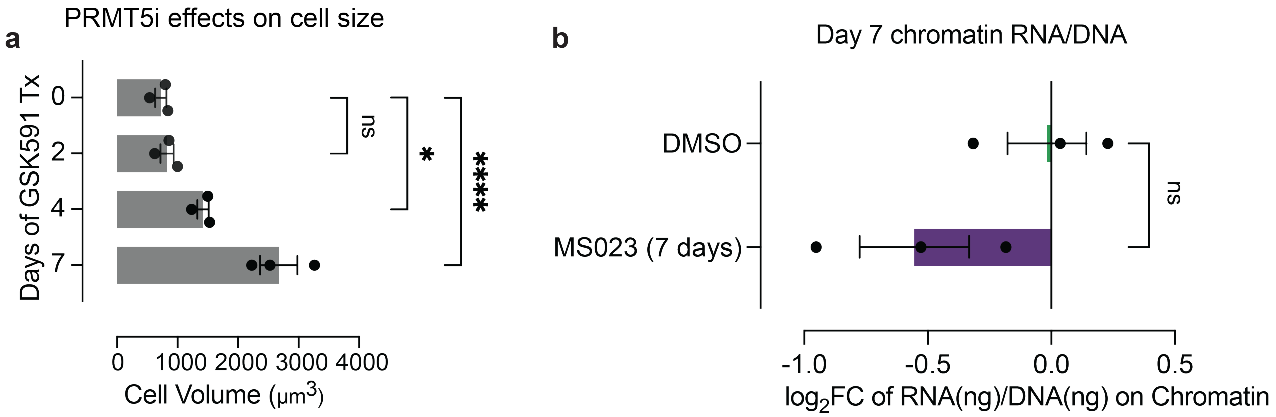


**Supplemental Figure S4 (Related to Figure 5). a)** Histogram depicting cell volume as a function of PRMT5 inhibition over time. *p<0.05, ****p<0.0001 **b)** Histogram of the $\log_2$ fold change of total chromatin RNA to DNA following Type I PRMT inhibition with MS023.

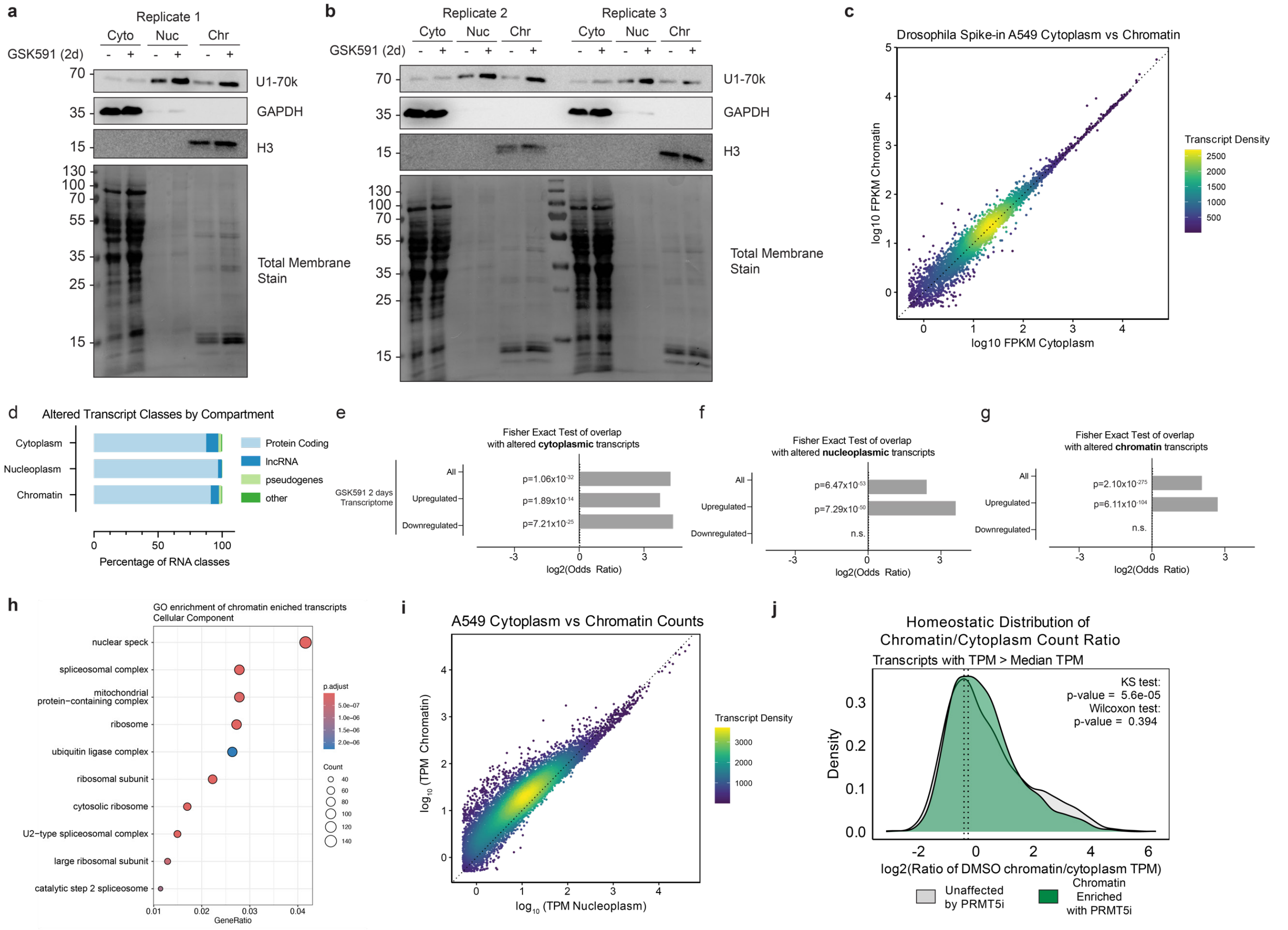


**Supplemental Figure S5 (Related to Figure 6). a-b)** Western blot controls of fractionated samples and replicates used for mRNA-sequencing. **c)** Correlation of the $\log_{10}$ average normalized read counts in FPKM for *D. melanogaster* transcripts in the cytoplasm and chromatin compartments for untreated cells (DMSO). Dotted line represents y=x. Transcript color represents the density of points on the plot. **d)** Relative percentages of RNA classes of significantly altered transcripts (padj<0.05) between DMSO and GSK591 treated cellular compartments. **e-g)** Fisher exact tests of significantly altered transcripts in cellular compartment compared to gene sets from total GSK591 treated mRNA sequencing. **h)** Dot plot of cellular component gene ontology for chromatin-enriched transcripts. Dot size is representative to the number of genes per category and color represents the p-adjusted value. **i)** Correlation of the $\log_{10}$ average normalized read counts in TPM for transcripts in the nucleoplasm and chromatin compartments for untreated cells (DMSO). **j)** Density plots of the ratio of normalized TPM counts per gene (with TPM > median of all expressed genes) between chromatin and cytoplasm compartments, comparing genes found to be enriched on chromatin upon PRMTi. Kolmogorov-Smirnov and Wilcoxon ranked sum tests were used to compare distributions.

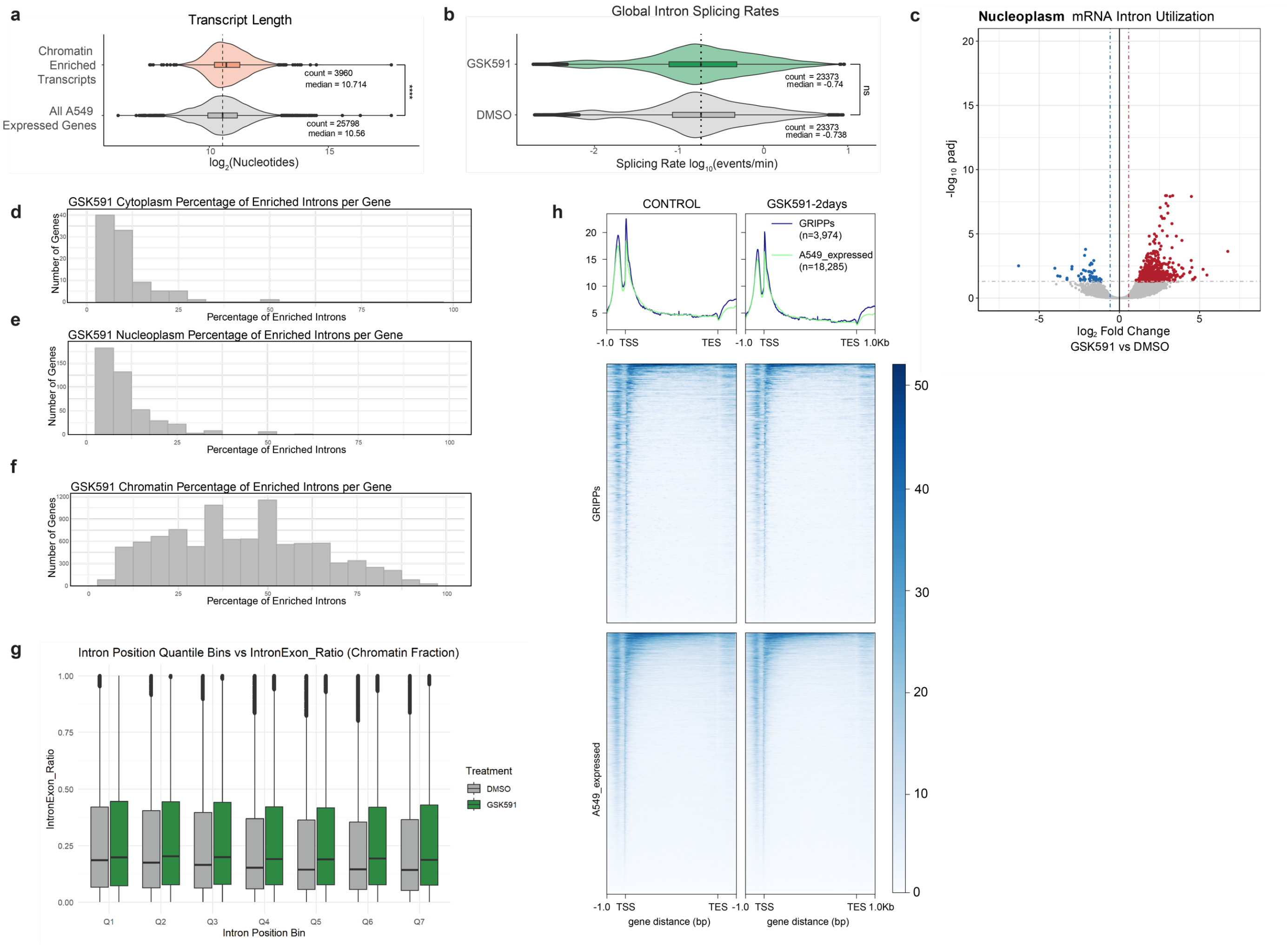


**Supplemental Figure S6 (Related to Figure 7). a)** Violin plots comparing transcript length of chromatin enriched genes upon PRMTi compared to all expressed genes in A549 cells. ****:p<0.0001 using Wilcoxon Ranked Sum test **b)** Splicing rates solved by SKaTER-seq comparing intron splicing rates solved both in GSK591 and DMSO treated conditions. Compared using Wilcoxon Ranked Sum test. **c)** Volcano plot of intron utilization for transcripts in nucleoplasm fraction comparing GSK591 vs DMSO. Red introns have a $log_2FC>0.58$ and $p_{adj}<0.05$. Blue introns have a $log_2FC<-0.58$ and a $p_{adj}<0.05$. **d-f)** Histograms depicting the percentage of enriched introns and gene number in GSK591 treated fractions. **g)** Intron:Exon Ratio for DMSO vs GSK591 Chromatin fraction across 7 quantiles throughout the transcripts. **h)** Pro-seq metagene profiles comparing GRIPPs to all genes expressed in A549 cells.

**Supplemental Table S5.** Antibodies used in this study

| Antibody Target | Vendor | Source | Identifiers | Additional Information |
|---|---|---|---|---|
| SNRPB | Proteintech | Rabbit polyclonal | Cat#: 16807-1-AP | WB: 1:2,000 |
| SNRPD3 | Abcam | Rabbit polyclonal | Cat#: ab157118 | WB: 1:2,000 |
| H3K27me3 | CST | Rabbit monoclonal | Cat#: 9733 | WB: 1:100,000 |
| Rme2s | Courtesy of Mark Bedford | Rabbit polyclonal | 4688 | WB: 1:10,000 |
| Rme2a | Cell Signaling | Rabbit monoclonal | Cat#: 8015S | WB: 1:2,000 |
| Rme1 | Cell Signaling | Rabbit monoclonal | Cat# 13522S | WB: 1:2,000 |
| H3 | Abcam | Rabbit polyclonal | Cat#: 1791 | WB: 1:300,000 |
| PRMT5 | Millipore | Rabbit polyclonal | Cat#: 07-405 | WB: 1:5,000 |
| MEP50 | LPBio | Rabbit polyclonal | Cat#: AR-0145-S | WB: 1:5,000 |
| pICln | Bethyl | Rabbit polyclonal | Cat#: A304-521A | WB: 1:50,000 |
| RIOK1 | Proteintech | Rabbit polyclonal | Cat#: 17222-1-AP | WB: 1:10,000 |
| GAPDH | Abcam | Mouse monoclonal | Cat#: ab9484 | WB: 1:50,000 |
| U1-snRNP 70 (E-4) | Santa Cruz | Mouse monoclonal | Cat#: SC390988 | WB: 1:2,000 |
| SNRPE | Proteintech | Rabbit polyclonal | Cat#: 20407-1-AP | WB:1:10,000 |
| IgG | Abcam | Rabbit polyclonal | Cat#: ab46540 | IP: 5 ug |
| m3G/TMG | MBL Life Science | Mouse monoclonal | Cat#: RN019M | IP: 5 ug |
| Rme2s | Cell Signaling | Rabbit monoclonal | Cat#: 13222S | WB: 1:2,000 |
| TBP | Cell Signaling | Rabbit polyclonal | Cat#: 8515 | WB: 1:10,000 |
| HA | Cell Signaling | Rabbit monoclonal | Cat#: C29F4 | WB: 1:100,000 |
| Coilin | BD Biosciences | Mouse monoclonal | Cat#: 612074 | WB: 1:250 |
| Tubulin (E7) | DSHB Iowa | Mouse monoclonal | Cat#: AB_2315513 | WB: 1:200 |

**Supplemental Table S6.** sgRNAs used in this study

| Target Gene | Guide # | Sequence |
|---|---|---|
| Scrambled Control | 1 | GTGTAGTTCGACCATTCGTG |
| PRMT5 | 3 | GGTCCCTCCCGCTGGACACG |
| PRMT5 | 4 | GAGAAAGATGGCGGCGATGG |
| COPRS (CoPR5) | 1 | GCCTGAAGGTCCATGCCCTG |
| COPRS (CoPR5) | 3 | CATGGACCTTCAGGCCGCCG |
| CLNS1A (pICln) | 2 | GCAGCAGAGTGCGGCAACAC |
| CLNS1A (pICln) | 3 | GCTGTGCTCCAACTCCCTCA |
| RIOK1 | 1 | AACCATTCAGAAGCCAACAG |
| RIOK1 | 3 | TGGCAGGGTGGTGGATCTGT |
| Coilin | 2 | GCCAGCTACCCCGCACTGTA |
| Coilin | 4 | CAAGATGGCAGCTTCCGAGA |
